# Redox Regulation of m^6^A Methyltransferase METTL3 in Human β-cells Controls the Innate Immune Response in Type 1 Diabetes

**DOI:** 10.1101/2023.02.16.528701

**Authors:** Dario F. De Jesus, Zijie Zhang, Natalie K. Brown, Xiaolu Li, Matthew J. Gaffrey, Sevim Kahraman, Jiangbo Wei, Jiang Hu, Giorgio Basile, Ling Xiao, Tariq M. Rana, Clayton Mathews, Alvin C. Powers, Mark A. Atkinson, Decio L. Eizirik, Sirano Dhe-Paganon, Audrey V. Parent, Wei-Jun Qian, Chuan He, Rohit N. Kulkarni

**Affiliations:** Section of Islet Cell and Regenerative Biology, Joslin Diabetes Center; Beth Israel Deaconess Medical Center; Harvard Medical School, Boston, USA; Department of Chemistry, Department of Biochemistry and Molecular Biology, and Institute for Biophysical Dynamics, The University of Chicago, Chicago, USA; Howard Hughes Medical Institute, The University of Chicago, Chicago, USA; Biological Sciences Division, Pacific Northwest National Laboratory, Richland, USA; Institute for Genomic Medicine, University of California, San Diego, La Jolla, USA; Department of Pathology, The University of Florida College of Medicine, Gainesville, USA; Department of Medicine, and Department of Molecular Physiology and Biophysics, Vanderbilt University Medical Center, Nashville, USA; ULB Center for Diabetes Research, Medical Faculty, Université Libre de Bruxelles (ULB), Brussels, Belgium; Department of Biological Chemistry, and Molecular Pharmacology, Harvard Medical School, Boston, USA; Diabetes Center, Department of Medicine, University of California, San Francisco, San Francisco, USA

## Abstract

Type 1 Diabetes (T1D) is characterized by autoimmune-mediated destruction of insulin-producing β-cells. Several observations have renewed interest in the innate immune system as an initiator of the disease process against β-cells. Here, we show that N^6^-Methyladenosine (m^6^A) is an adaptive β-cell safeguard mechanism that accelerates mRNA decay of the 2’-5’-oligoadenylate synthetase (OAS) genes to control the antiviral innate immune response at T1D onset. m^6^A writer methyltransferase 3 (METTL3) levels increase drastically in human and mouse β-cells at T1D onset but rapidly decline with disease progression. Treatment of human islets and EndoC-βH1 cells with pro-inflammatory cytokines interleukin-1 β and interferon α mimicked the METTL3 upregulation seen at T1D onset. Furthermore, m^6^A-sequencing revealed the m^6^A hypermethylation of several key innate immune mediators including *OAS1, OAS2,* and *OAS3* in human islets and EndoC-βH1 cells challenged with cytokines. METTL3 silencing in human pseudoislets or EndoC-βH1 cells enhanced OAS levels by increasing its mRNA stability upon cytokine challenge. Consistently, *in vivo* gene therapy, to prolong Mettl3 overexpression specifically in β-cells, delayed diabetes progression in the non-obese diabetic (NOD) mouse model of T1D by limiting the upregulation of *Oas* pointing to potential therapeutic relevance. Mechanistically, the accumulation of reactive oxygen species blocked METTL3 upregulation in response to cytokines, while physiological levels of nitric oxide promoted its expression in human islets. Furthermore, for the first time to our knowledge, we show that the cysteines in position C276 and C326 in the zinc finger domain of the METTL3 protein are sensitive to S-nitrosylation (SNO) and are significant for the METTL3 mediated regulation of OAS mRNA stability in human β-cells in response to cytokines. Collectively, we report that m^6^A regulates human and mouse β-cells to control the innate immune response during the onset of T1D and propose targeting METTL3 to prevent β-cell death in T1D.

## INTRODUCTION

Type 1 Diabetes (T1D) is characterized by immune cell infiltration of pancreatic islets which leads to the selective destruction of β-cells. During the autoimmune attack, the release of cytokines by immune mediators utilizes complex pathways to promote β-cell dysfunction and apoptosis (Atkinson et al., 2015; Coomans de Brachène et al., 2018). The precise mechanisms that trigger the cascade of events that ultimately lead to β-cell death are not fully understood. Physiological β-cell death at early stages of development has been proposed as a mechanism triggering self-reactive T-cells (Turley et al., 2003). Recent findings suggest the ability of metabolic abnormalities to trigger stress pathways that promote the deterioration of β-cell function prior to seroconversion in T1D (Sims et al., 2020). These studies have prompted a renewed interest in the central role of the β-cell as a target cell that initiates the disease process (Roep et al., 2021; Sims *et al.,* 2020; Wilcox et al., 2016).

The ability to distinguish self from non-self DNA or RNA is a fundamental function of the innate immune system. For example, immune responses to viral infections are almost entirely dependent on the activation of innate immune sensors that are able to systematically survey cells for the presence of foreign nucleic acids (Crowl et al., 2017). Consequently, several autoimmune diseases are triggered by the over-activation of the innate immune system (Lang et al., 2007).

Innate immunity is genetically linked with T1D, with several GWAS signals associated with innate immune mediators (Bluestone et al., 2010). Furthermore, ample evidence points to activation of the innate immune system during early stages of T1D. For example, several innate immune mediators including interferon genes are upregulated prior to the onset of T1D (Carry et al., 2022; Kallionpaa et al., 2014). Additionally, multiple nucleic acid sensors involved in the innate immune response are upregulated in insulitic islets at T1D onset (Apaolaza et al., 2021; Lundberg et al., 2016). Among the upregulated genes include the oligoadenylate synthase (OAS) family of proteins, a class of nucleotidyltransferases that, once activated by type I interferon or double-stranded RNA (dsRNA), either act independently or produce 2ū–5ū-linked oligoadenylates to activate RNase L (Hornung et al., 2014). Sustained activation of OAS or RNAse L may lead to protein translation arrest and cell apoptosis (Hornung *et al.,* 2014). Indeed, polymorphisms in the OAS gene cluster have been associated with increased OAS enzymatic activity and susceptibility to T1D (Field et al., 2005; Pedersen et al., 2021; Tessier et al., 2006). Intriguingly, β-cells are unique among pancreatic islet cells with the ability to upregulate OAS expression in response to interferon-α or poly(I:C) (a dsRNA mimetic) (Bonnevie-Nielsen et al., 1996; Li et al., 2009b). OAS overexpression in β-cells leads to proliferation arrest and apoptosis (Dan et al., 2012; Li *et al.,* 2009b), while mice deficient in RNase L are protected from diabetes in a dsRNA-induced mouse model of T1D, consistent with the notion that over-activation of the OAS - RNase L pathway leads to β-cell death and T1D (Zeng et al., 2014).

N6-methyladenosine (m^6^A) is the most abundant modification in mRNA in virtually all mammals (Dominissini et al., 2012; Frye et al., 2018; Meyer et al., 2012). Adenosine methylation levels are regulated by protein complexes, including “writers” such as methyltransferase 3 (METTL3) and 14 (METTL14) (Frye *et al.,* 2018). Several RNA binding proteins – “readers”, including YT521-B homology family proteins (e.g.,YTHDF1, YTHDF2, and YTHDF3), recognize methylated adenosines and regulate several aspects of mRNA biology including mRNA decay (Lasman et al., 2020; Lee et al., 2021; Zou et al., 2022). METTL3 is the only enzyme in the m^6^A writer complex that presents catalytic activity (Wang et al., 2016). Furthermore, the consequences of activation and post-translational modification of METTL3 are not fully explored. For example, recent work has demonstrated that METTL3 activity can be regulated by SUMOylation (Du et al., 2018) and phosphorylation (Sun et al., 2020). However, the role of cysteine oxidative modifications such as S-nitrosylation (SNO) has not been explored.

Recent discoveries have proposed that the m^6^A machinery can modify viral as well as cell host mRNAs to impact infection outcomes (McFadden and Horner, 2021). More specifically, m^6^A has been shown to suppress the innate immune response by accelerating the turnover of type I interferon genes in fibroblasts (Winkler et al., 2019) by promoting adenosine-to-inosine (A-to-I) RNA editing through regulation of ADAR1 (Terajima et al., 2021) or by blocking the synthesis of endogenous aberrant dsRNAs (Gao et al., 2020; Qiu et al., 2021). However, the biological roles of m^6^A in T1D and more specifically, their contribution towards mediating β-cell innate immune responses are virtually unknown.

Here, we characterize for the first time to our knowledge, the contribution of m^6^A to the pathophysiology of T1D. First, a careful examination of the expression of the m^6^A modulators in mouse or human T1D β-cells revealed an upregulation of METTL3 at the onset of T1D followed by a rapid decline as the disease progressed. Next, to phenocopy the alterations in m^6^A levels during the onset of T1D we used an *in vitro* approach by treating human islets or EndoC-βH1 cells with a combination of interferon α (IFN-α) and interleukin-1 β (IL-1β) followed by mapping the β-cell m^6^A landscape using m^6^A-seq and RNA-seq. Consistent with our hypothesis, we identified m^6^A hypermethylation of OAS genes and demonstrated that METTL3 downregulation in human pseudoislets or EndoC-βH1 cells leads to the upregulation of OAS proteins. Furthermore, we demonstrated that m^6^A accelerates the mRNA decay of OAS via S-nitrosylation of the cysteine residues (C276 and C326) in the redox-sensitive zinc finger domain of METTL3. The ability of sustained over-expression of Mettl3 in β-cells to limit the up-regulation of Oas and protect NOD mice from developing diabetes supports the translational significance of these findings.

Together, our studies identify m^6^A as a novel adaptive β-cell safeguard mechanism that controls the innate immune response at T1D onset and reveals S-nitrosylation as a novel post-translational modification that controls METTL3 function. Our results point to targeting the m^6^A writer METTL3 to promote β-cell survival with the long-term goal of counteracting hyperglycemia.

## RESULTS

### m^6^A writer (METTL3) levels peak at the onset and decrease drastically with the progression of T1D

To begin an exploration into the role of m^6^A in immune-mediated events associated with β-cell loss in Type 1 Diabetes (T1D) we re-analyzed an RNA-seq dataset performed in sorted β-cells from Mettl14 KO and control mice (De Jesus et al., 2019). Specifically, we performed pathway analyses on the upregulated genes (Figure S1A; Figure S1B) and observed enrichment in pathways associated with the immune response, including complement activation, antigen processing and presentation, and the innate immune system (Figure S1B). Among the immune system-related upregulated genes in β-cells harboring Mettl14 KO were several genes associated with the immunogenetics of T1D including the histocompatibility 2, class II antigen A, alpha *(H2-Aa),* CD74 molecule *(Cd74),* and CD44 molecule *(Cd44)* (Figure S1C).

Next, to determine if the m*^6^*A modulators, and in particular the main m^6^A writers, are impacted by T1D, we isolated islets from the non-obese diabetic (NOD) mouse model of T1D (Makino et al., 1980) and from a NOD congenic insulitis and diabetes resistant (NOR) model that was used as a near genetically identical control (Prochazka et al., 1992). Gene expression analyses revealed an upregulation of *Mettl3* and *Mettl14* in islets from NOD compared to NOR mice at 4 weeks of age (Figure 1A), followed by their downregulation at age 8 weeks (Figure 1A). To characterize the transcriptomic profile and to confirm whether the changes in the expression of the m^6^A writers were specific to β-cells, we initially performed fluorescence-activated cell sorting (FACS) of isolated and dispersed NOD islets from 4- and 8-week-old mice. The β- and non-β-cells were sorted by autofluorescence (Smelt et al., 2008), negatively selected for CD45 (a marker of hematopoietic cells) (Figure S2A) and subjected to RNA-seq. The CD45 negatively enriched β-cell and non-β-cell populations segregated by group (Figure S2B), with the β-cell fractions enriched for β-cell identity genes compared to non-β-cell fractions (Figure S2C). Interestingly, β-cells from 8-week old NOD mice presented upregulation of several genes involved in T1D (Figure S2D), including antigen processing and presentation (Figure S2E), and innate immunity (Figure S2F). Gene expression analyses of m^6^A writers revealed a tendency to decrease in *Mettl3* while a significant downregulation was evident in *Mettl14* and *Wtap* specifically in β-cell fractions in the 8-week old compared to the 4-week old NODs (Figure 1B).

**Figure 1:**
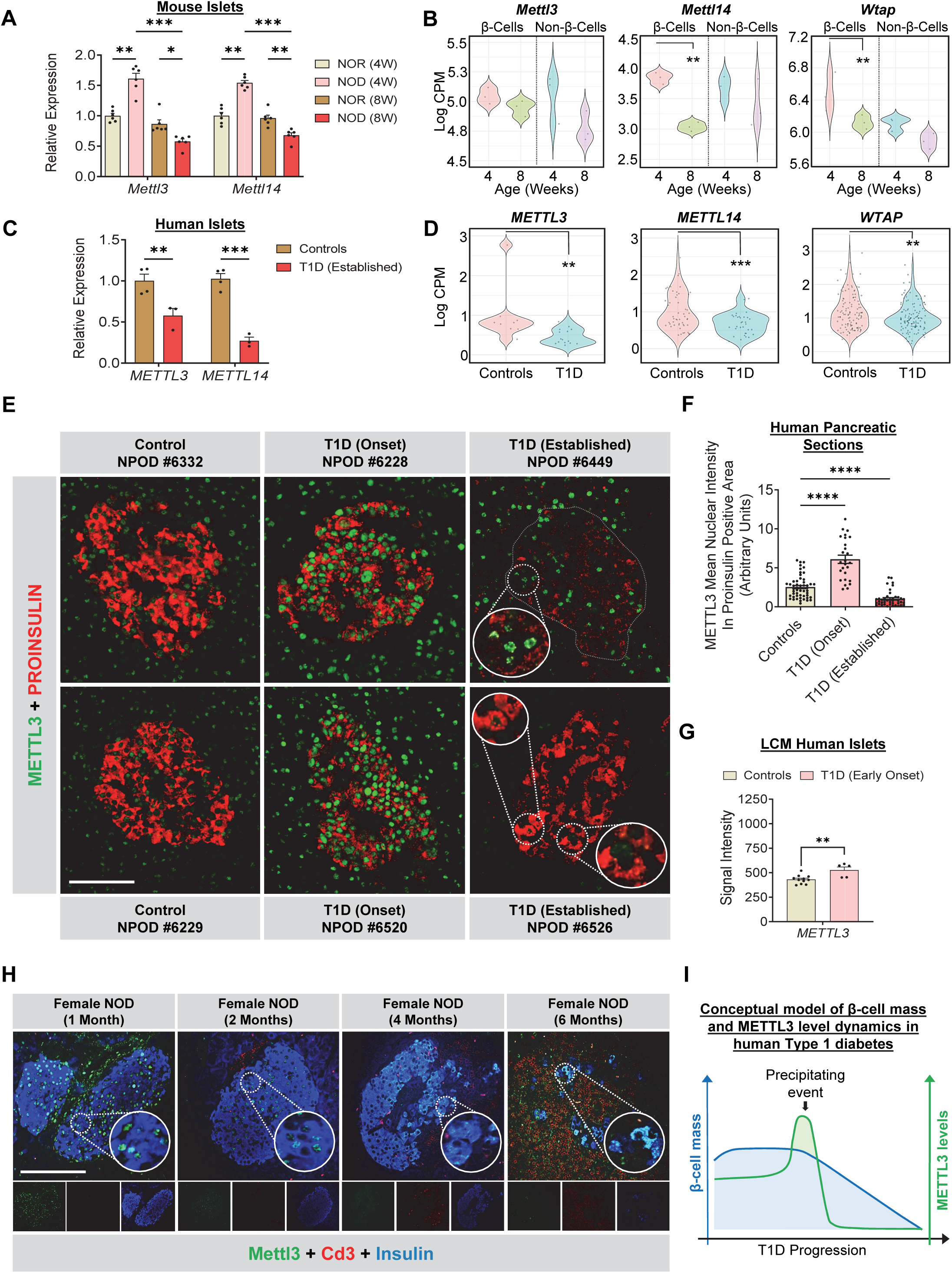
m^6^A writer METTL3 levels peak at Type 1 diabetes onset but decrease drastically with disease progression. **(A)** qRT-PCR analyses of the m^6^A writer’s genes in whole islets isolated from 4- week-old or 8-week-old NOR or NOD mice (n=6 mice/group). **(B)** Violin-plots representation of the distribution of gene expression of m^6^A writers in sorted β- and non-β-cells in 4- or 8-week-old NOD mice (n=3 pools of 3 mice/group). **(C)** qRT-PCR analyses of *METTL3* and *METTL14* in whole islets isolated from human non-diabetic Controls (n=4), and human patients with established T1D (n=3 samples). **(D)** Violin-plots representation of the distribution of gene expression of m^6^A writers in single β-cells (high insulin gene expression) from Control (n=12) and established T1D (n=4) (GSE121863). **(E)** Representative pictures of immunofluorescence staining METTL3 in pancreatic sections collected from non-diabetic (Control) humans (n=9), T1D Onset (n=4), and established T1D (n=7) (Scale bar = 100 µm). **(F)** Quantification of METTL3 intensity in proinsulin positive area in pancreatic sections collected from non-diabetic (Control) humans (n=9), T1D Onset (n=4), and established T1D (n=7). **(G)** Microarray signal intensity of *METTL3* in laser-capture microdissected (LCM) pancreatic islets of non-diabetic (Control) humans (n=11), and T1D Onset insulitic islets (n=5). **(H)** Representative pictures of immunofluorescence staining of METTL3 in pancreatic sections collected from NOD female mice with 1-, 2-, 4, and 6 months of age (n=3) (Scale bar = 50µm). **(I)** Conceptual schematic representation of METTL3 and β-cell mass dynamics in the progression of humanT1D. All samples in each panel are biologically independent. Data were expressed as means ± SEM. \**P*<0.01, \*\**P*<0.01, \*\*\**P*<0.001. Statistical analysis was performed by Two-Way ANOVA with Turkey multiple comparison test in A. Benjamini-Hochberg procedure in B and D. Two-Way ANOVA with Holm-Sidak’s multiple comparisons test in C, and F. Two-tailed unpaired t-test in G.

T o further examine the impact of T1D on the expression of the m^6^A writers, we obtained freshly isolated human islets from patients with established T1D or non-diabetic controls (patient information in Supplementary Table 1) and cultured them overnight before subjecting them to qPCR analyses. Gene expression of *METTL3* and *METTL14 was* dramatically downregulated in established T1D islets compared to controls (Figure 1C). To confirm this is a consistent finding we downloaded and re-analyzed a single-cell RNA-seq dataset performed in islets from an independent cohort of established T1D patients and non-diabetic controls (Russell et al., 2019). Pancreatic β-cells were identified by insulin *(INS)* gene expression (Figure S3A) and easily segregated from other islet cell types including α-cells expressing glucagon *(GCG)* (Figure S3B). The expression of the m^6^A writers *METTL3, METTL14,* and *WTAP* were downregulated in β-cells from patients with established T1D compared to non-diabetic controls (Figure 1D). Overall, these observations suggest that m^6^A writers are downregulated in established T1D in mice and humans.

METTL3 is considered an attractive therapeutic target considering it is the only subunit of the m^6^A writer complex presenting enzymatic activity (Sledz and Jinek, 2016; Wang *et al.,* 2016). Consequently, recent research efforts have focused on identifying potent and selective METTL3 catalytic inhibitors to counter pathological states such as leukemia (Yankova et al., 2021) and renal injury (Wang et al., 2022). We took advantage of the abundance of METTL3 in human β-cells (De Jesus *et al.,* 2019) to test the dynamic regulation of its expression at various stages of T1D development. Specifically, we performed immunofluorescence staining of METTL3 and proinsulin (as a β-cell marker) in human pancreatic sections from control, T1D onset (at the onset of the disease at hospital admission), or established T1D (nPOD patient information in Supplementary Table 1). Using a pipeline on ImageJ (Schneider et al., 2012) to unbiasedly measure METTL3 nuclear intensity in the proinsulin positive area (Figure S3C) across different groups, we detected an upregulation of METTL3 across all T1D onset samples compared to controls, followed by a downregulation in established T1D cases (Figure 1E and F). METTL3 upregulation was also noticeable in the acinar tissue at T1D onset (Figure 1E). The ease of identification of human islets from T1D onset patients even without proinsulin co-staining reflected the robust upregulation of METTL3 predominantly in islets and specifically in β-cells (Figure S3D). Re-analyses of a microarray dataset (Gerling, Mathews et al; unpublished) performed in laser-capture micro-dissected human islets exhibiting insulitis at T1D onset versus insulitis-free control islets confirmed the upregulation of METTL3 in the former and validated our observations (Figure 1G).

T1D is a heterogeneous disease (Insel et al., 2015). While nearly 70% of the children at genetic risk who present multiple islet autoantibodies eventually develop T1D, the progression to overt disease following seroconversion may take decades (Ziegler et al., 2013). In contrast, 19% of the patients develop T1D without seroconversion (Wang et al., 2007). Consequently, mapping temporal changes using the available pancreases from patients with T1D, for pragmatic and ethical reasons, continues to be a challenge and prompted the use of animal models. Therefore, to further dissect the METTL3 dynamics in T1D progression, we collected pancreatic sections from NOD mice at stages of prediabetes, diabetes, or overt diabetes for immunostaining for Mettl3, insulin, and Cd3 (a T cell marker) (Figure 1H and Figure S3E). Consistent with the human data, protein levels of Mettl3 decreased in female NODs with T1D progression (Figure 1H), while being unaltered in age-matched males (which are known to not develop insulitis or diabetes) (Figure S3E).

Altogether these data demonstrate the existence of a conserved and dynamic regulation of METTL3 during progression of T1D in mice and humans. Early onset of the disease is characterized by a significant upregulation of METTL3 levels in β-cells followed by its downregulation that spans established T1D (Figure 1I) and coincides with the notion of a “precipitating event” as postulated by Eisenbarth (Eisenbarth, 1986).

### Stimulation of human β-cells with IL-1β and IFN-α recapitulates the upregulation of METTL3 observed at the onset of T1D

Next, we aimed to create an *in vitro* tool to recapitulate and study the METTL3 upregulation seen at T1D onset in human β-cells. The early inflammation associated with the accumulation of immune cells during the initial stages of T1D is thought to be mediated by inflammatory cytokines such as interleukin-1β (IL-1β), interferon α (IFN-α), and interferon γ (IFN-/) (Eizirik et al., 2009; Todd, 2010). Cytokine treatment of human islets, and in particular β-cells, has been reported to recapitulate several pathophysiological aspects of the molecular landscape of T1D (Colli et al., 2018; Eizirik et al., 2012; Ramos-Rodriguez et al., 2019).

To test whether proinflammatory cytokines impact m^6^A levels and METTL3 in particular, we challenged human islets or a human β-cell line (EndoC-βH1) (Benazra et al., 2015) with IL-1 β, IFN-α, IFN-γ, or a combination of IL-1 β + IFN-α, or IL-1 β + IFN-γ (Figure 2A) for 48h. We then measured total m^6^A levels by LC-MS/MS (see methods) and analyzed METTL3 and METTL14 protein levels (Figure 2A). m^6^A levels were upregulated by IFN-α and even more robustly by the combination of IL-1 β + IFN-α (Figure 2B and C). Overall, METTL3 and METTL14 protein levels increased significantly in human islets in response to treatment with proinflammatory cytokines (Figure 2D). On the other hand, EndoC-βH1 cells exhibited upregulated METTL3 specifically upon the addition of IL-1β, IFN-α, or a combination of IL-1 β + IFN-α (Figure 2E). Together, these results indicate that stimulation of human β-cells with IL-1 β + IFN-α leads to the upregulation of METTL3 and m^6^A levels recapitulating T1D onset. Next, to further explore METTL3 upregulation dynamics, we challenged human islets with IL-1 β + IFN-α or PBS for 24-, 48-, or 72h (Figure 2F). METTL3 and METTL14 showed a graded increase with stimulation time, with METTL14 showing a significant increase after 48h of stimulation with IL-1 β and IFN-α (Figure 2G). Finally, to visually confirm METTL3 upregulation in response to cytokines, we challenged human islets with IL-1 β and IFN-α for 48h and performed immunofluorescence imaging on agar-embedded islets (Figure 2H). As expected, METTL3 levels were increased upon IL-1 β and IFN-α stimulation (Figure 2H), with said upregulation occurring largely in insulin-positive islet cells. Overall, these results demonstrate that stimulating human islets and β-cells with IL-1β and IFN-α *in vitro* recapitulates the upregulation of METTL3 seen at human T1D onset.

**Figure 2:**
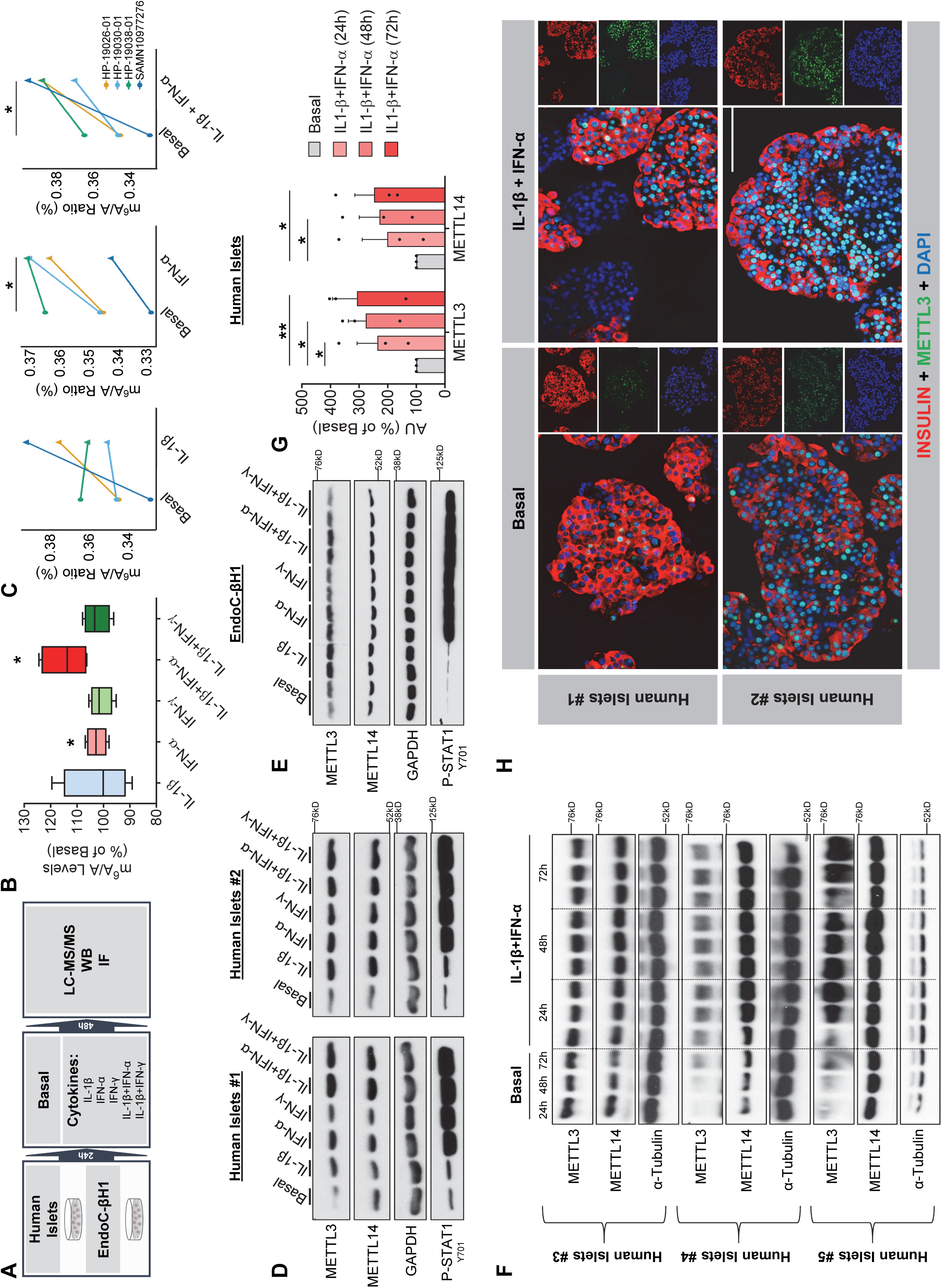
Co-treatment of human β-cells with Interleukin 1 beta (IL-1 β) and interferon alpha (IFN-α) recapitulates the METTL3 upregulation seen at human T1D onset. **(A)** Summary scheme of the experimental plan. **(B)** m^6^A levels measured by LC-MS/MS in human islets treated with IL-1 β, IFN-α, interferon-gamma (IFN-γ), a combination of IL-1 β plus IFN-α, or IL1-β plus IFN-γ for 48h compared to PBS-treated (n=4). **(C)** m^6^A levels measured by LC-MS/MS in individual human islet donors treated with Il1-β, IFN-α, or IL-1 β plus IFN-α compared to basal (PBS-treated) (n=4). Box plot shows the median, box edges show first and third quartiles, and whiskers show the minimum and maximum. **(D)** Western-blot analyses of indicated proteins in human islets treated with the represented cytokines or cytokine combinations for 48h (n=2). **(E)** Western-blot analyses of indicated proteins in EndoC-βH1 cells treated with the represented cytokines or cytokine combinations for 48h (n=2). **(F)** Western-blot analyses of indicated proteins in human islets treated with IL-1 β plus IFN-α or PBS (Basal) for 24-, 48-, or 72h (n=3). **(G)** Protein quantification of (F). **(H)** Representative pictures of immunofluorescence staining analyses of METTL3 in agar-embedded islets collected from human islets treated with IL-1 β plus IFN-α or PBS (Basal) for 48h (n=2) (Scale bar =100 µm). All samples in each panel are biologically independent. Data were expressed as means ± SEM. \**P*<0.01, \*\**P*<0.01. Statistical analysis was performed by two-tailed paired t-test in B and C. Two-Way ANOVA with Fisher’s LSD test in G.

### m^6^A landscape analyses of human islets treated with IL-1 β and IFN-α reveal hypermethylation of antiviral innate 2’,5’-oligoadenylate synthetase (OAS) genes

To directly evaluate the impact of the dynamic expression of METTL3 on the m^6^A landscape during T1D onset, we employed RNA-seq and m^6^A-seq in islets from 15 independent human donors (Supplementary Table 1) stimulated with a combination of cytokines (IL-1 β+IFN-α) or PBS for 48h (Figure 3A). The transcriptome (Figure 3B) and the m^6^A methylome (Figure 3C) of (IL-1 β+IFN-α)-treated islets showed segregation from PBS-treated samples. We began by studying the transcriptomic changes induced by cytokines and observed downregulation of 5181 genes and the upregulation of 5005 genes compared to PBS-treated samples (FDR<0.05) (Figure 3D). This suggested the ability of IL-1β and IFN-α to induce significant transcriptomic remodeling in human islets. Enriched pathway analyses of the combined upregulated and downregulated gene sets (FDR<0.05) revealed an enrichment in pathways involved in nonsense-mediated decay and several innate immune pathways including interferon signaling, ISG15 antiviral mechanism, innate immune response to cytosolic DNA, and RIG-receptor signaling (Figure 3E).

**Figure 3:**
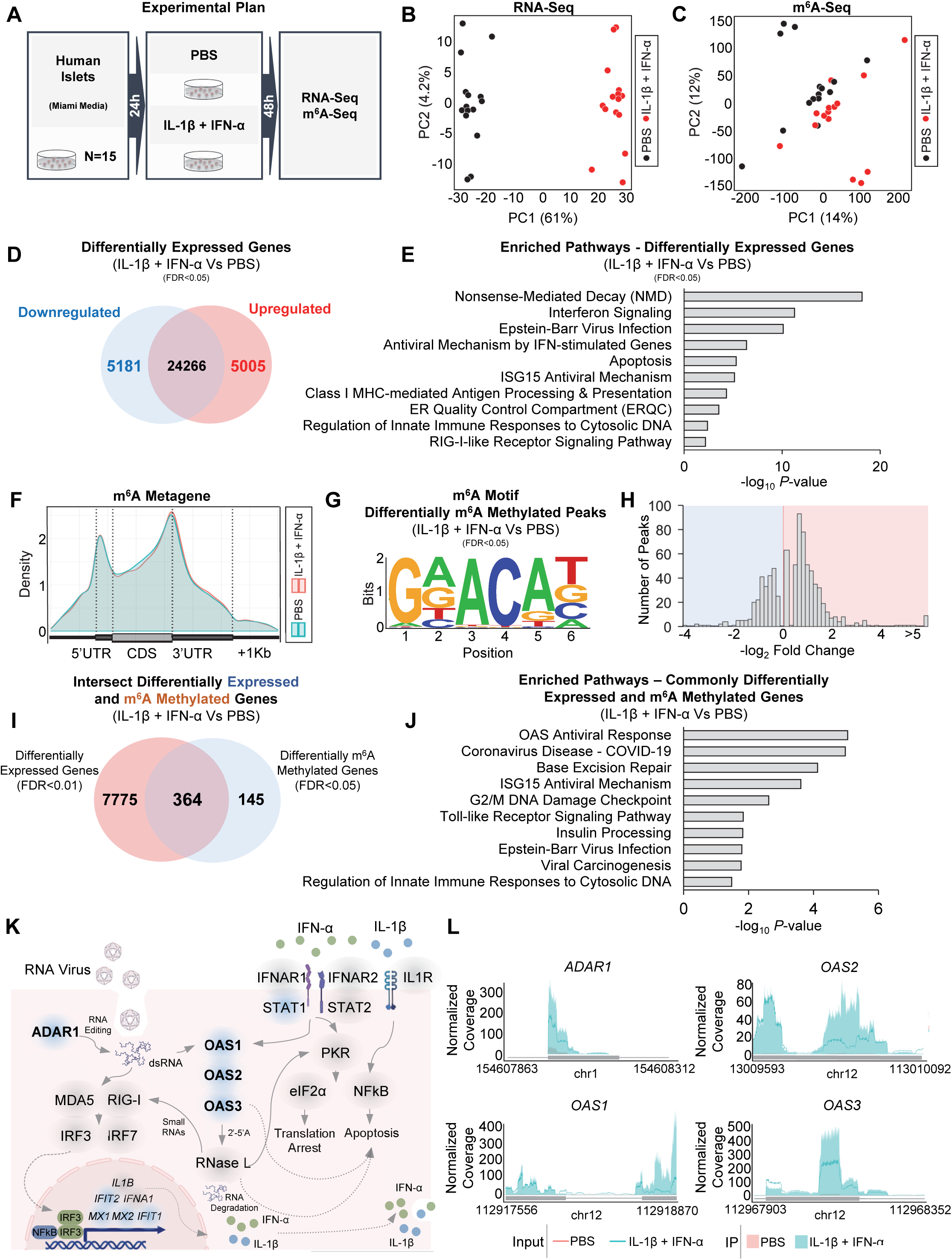
m^6^A landscape analyses of human islets treated with IL-1β and IFN-α reveal hypermethylation of 2’,5’-oligoadenylate synthetase (OAS) genes. **(A)** Summary scheme of the experimental plan. **(B)** PCA plot of RNA-sequencing in human islets treated with PBS (black dots) or IL-1β and IFN-α (red dots) (n=15). **(C)** PCA plot of m^6^A-sequencing in human islets treated with PBS (black dots) or IL-1 β and IFN-α (red dots) (n=15). **(D)** Venn diagram representation of the upregulated (red), downregulated (blue), and unchanged genes (black) in human islets treated with IL-1 β and IFN-α compared to PBS. Statistical analyses were performed using the Benjamini-Hochberg procedure and genes were filtered for FDR<0.05. **(E)** Pathway enrichment analyses of upregulated and downregulated genes in human islets treated with IL-1β and IFN-α compared to PBS. *P*-values were calculated according to the hypergeometric test based on the number of physical entities present in both the predefined set and user-specified list of physical entities. **(F)** Metagene of m^6^A enriched peaks in PBS-(blue) and IL-1 β plus IFN-α-treated (red) human islets. **(G)** Enrichment for known m^6^A consensus motif RRACH. **(H)** Histogram of log_2_-fold change showing the distribution of differential m^6^A loci fold changes from IL-1β plus IFN-α-treated versus PBS. **(I)** Venn diagram representation of the intersection between differentially methylated and expressed genes in human islets treated with IL-1 β and IFN-α compared to PBS (n=15). Statistical analyses were performed using the Benjamini-Hochberg procedure and differentially expressed genes were filtered for FDR<0.01 and m^6^A methylated genes for FDR<0.05. **(J)** Pathway enrichment analyses of intersected genes in (I). **(K)** Representation of antiviral innate immune pathway based on KEEG and Wikipathway annotations depicting several m^6^A hypermethylated genes (blue shade) and unchanged genes (grey shade) in human islets treated with IL-1β and IFN-α compared to PBS-treated (genes filtered for FDR<0.05). **(L)** Coverage plots of m^6^A peaks in *ADAR1, OAS1,* OAS2, and *OAS3* genes in human Islets treated with L-1 β and IFN-α (blue) or PBS (red). Plotted coverages are the median of the n replicates presented. All samples in each panel are biologically independent.

Next, analyses of the m^6^A methylome of (IL-1 β+IFN-α)-treated human islets showed changes that were consistent with patterns from previous studies (Dominissini *et al.,* 2012; Meyer *et al.,* 2012) in that the m^6^A peaks were enriched at the start and stop codons (Figure 3F) and were characterized by the canonical GGACU motif (Figure 3G). Differential analysis of m^6^A-sequencing revealed 800 differently methylated sites in 509 genes (FDR<0.05) with a higher number of hypermethylated m^6^A sites in IL-1 β and IFN-α-treated compared to PBS-treated islets (Figure 3H). Since IL-1 β and IFN-α stimulation of human islets led to extensive transcriptomic remodeling, we hypothesized that some of these pathways are modulated by m^6^A. To test this possibility, we intersected the differentially expressed (DEGs) and m^6^A methylated genes (DMGs) in human islets treated with IL-15 and IFN-α compared to PBS. Pathway enrichment analyses on these common 364 intersected genes (Figure 3I) revealed several interconnected pathways involving the antiviral innate immunity system (Figure 3J). More specifically, several double-strand RNA sensors such as the 2’,5’-oligoadenylate synthetase (*OAS*) antiviral response genes were highly hypermethylated in human islets in response to IL-1β and IFN-α, highlighted by the genes in this pathway including *OAS1, OAS2,* and *OAS3* (Figure 3K and L). Other hypermethylated genes included adenosine deaminase acting on RNA *(ADAR1)* (Figure 3K and L) as well as several interferon-induced genes such as interferon-induced protein with tetratricopeptide *(IFIT1* and *IFIT2),* and MX dynamin-like GTPase *(MX1* and *MX2)* (Figure 3K). Overall, these results indicate a dynamic transcriptomic remodeling in human islets upon IL-1β and IFN-α stimulation and show that genes differentially expressed and m^6^A-decorated appear to be mainly involved in the antiviral innate immune response, particularly the 2’,5-oligoadenylate synthetase pathway.

### OAS genes are upregulated and m^6^A decorated in human β-cells treated with IL-1 β and IFN-α

To begin to validate the m^6^A regulation of OAS genes specifically in human β-cells, we challenged EndoC-βH1 cells (a pure human β-cell line) with (IL-1β+IFN-α) or PBS for 48h followed by RNA-seq and m^6^A-seq (Figure 4A). EndoC-βH1 is a human β-cell line expressing METTL3 (De Jesus *et al.,* 2019) and β-cell identity genes, and has been shown to respond to cytokines, and exhibit glucose-stimulated insulin secretion (Hastoy et al., 2018; Tsonkova et al., 2018).

**Figure 4:**
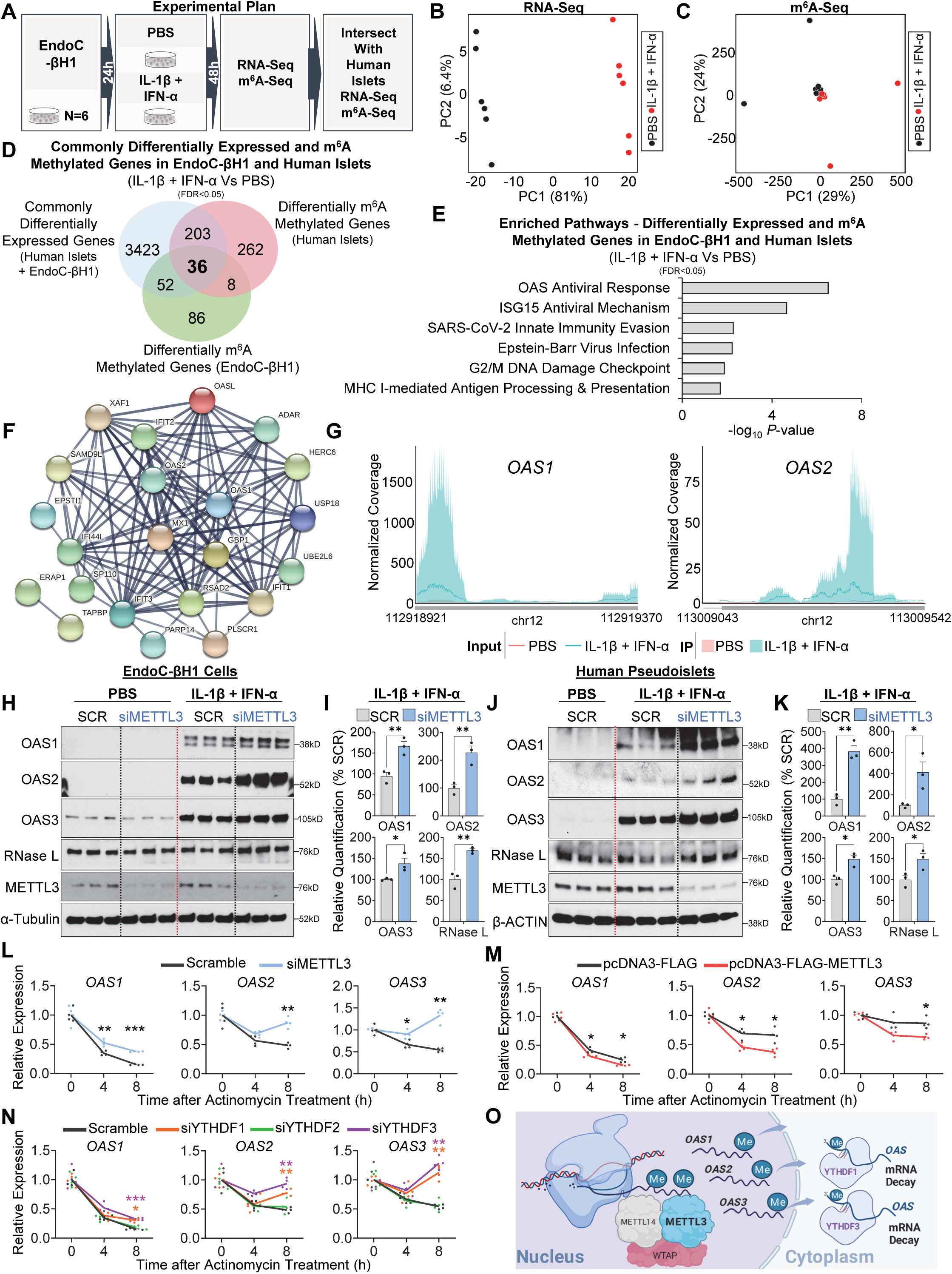
m^6^A controls the mRNA stability of 2’-5’-oligoadenylate synthetase (OAS) genes. **(A)** Summary scheme of experimental plan. **(B)** PCA plot of RNA-sequencing in EndoC-βH1 cells treated with PBS (black dots) or IL-1 β and IFN-α (red dots) (n=6). **(C)** PCA plot of m^6^A-sequencing in EndoC-βH1 treated with PBS (black dots) or IL-1β and IFN-α (red dots) (n=6). **(D)** Venn diagram representation of the intersection between differentially expressed genes in human islets and EndoC-βH1 treated with IL-1 β and IFN-α compared to PBS-treated with differentially m^6^A methylated genes in human islets and EndoC-βH1 treated with IL-1β and IFN-α compared to PBS-treated. Statistical analyses were performed using the Benjamini-Hochberg procedure and genes were filtered for FDR<0.05. **(E)** Pathway enrichment analyses of intersected genes in (D). *P*-values were calculated according to the hypergeometric test based on the number of physical entities present in both the predefined set and user-specified list of physical entities. **(F)** STRING functional protein-protein interaction network of 36 intersected genes, showing differential expression and m^6^A methylation in human islets and Endoc-βH1 cells treated with IL-1 β and IFN-α compared to PBS. **(G)** Coverage plots of m^6^A peaks in *OAS1* and *OAS2* genes in EndoC-βH1 cells treated with L-1 β and IFN-α or PBS. Plotted coverages are the median of the n replicates presented. **(H)** Western-blot analyses of indicated proteins after IL-1β plus IFN-α or PBS stimulation in EndoC-βH1 cells harboring METTL3 KD or scramble (SCR) (n=3). **(I)** Protein quantification of indicated protein in (H). **(J)** Western-blot analyses of indicated proteins after IL-1 β plus IFN-α or PBS stimulation in human pseudoislets METTL3 KD or SCR (n=3). **(K)** Protein quantification of indicated protein in (J). **(L)** qRT-PCR analyses of OAS genes after IL-1 β plus IFN-α stimulation of METTL3 KD or scramble EndoC-βH1 cells after a time-course treatment with actinomycin D (ActD) (n=4/group). **(M)** qRT-PCR analyses of OAS genes after IL-1 β plus IFN-α stimulation in METTL3 overexpressing (OE) and control (FLAG) EndoC-βH1 cells after a time-course treatment with ActD (n=4/group). **(N)** qRT-PCR analyses of OAS genes after IL-1β plus IFN-α stimulation in scramble, YTHDF1, YTHDF2, or YTHDF3 KD EndoC-βH1 cells after a time-course treatment with ActD (n=4/group). **(O)** Model depicting the role of METTL3-mediated m^6^A hypermethylation of OAS genes in response to IL-1 β and IFN-α or at T1D onset that leads to OAS nuclear export to the β-cell cytoplasm and recognition by m^6^A readers YTHDF1 and YTHDF3 leading to accelerated mRNA decay. All samples in each panel are biologically independent. Data were expressed as means ± SEM. \**P*<0.01, \*\**P*<0.01, \*\*\**P*<0.001. Statistical analysis was performed by two-tailed unpaired t-test or as otherwise stated above.

The transcriptome (Figure 4B) and the m^6^A methylome (Figure 4C) of IL-1β and IFN-α-treated versus PBS-treated EndoC-βH1 cells showed clear segregation on dimension-reduced data. IL-1β and IFN-α stimulation resulted in the downregulation of 3166 genes and the upregulation of 3203 genes compared to PBS-treated EndoC-βH1 cells (FDR<0.05) (Figure S4A). This suggests that similar to human islets, EndoC-βH1 cells exhibit notable transcriptomic remodeling in response to IL-1 and IFN-α. Pathway enrichment analyses of the combined upregulated and downregulated genes (FDR<0.05) revealed pathways involved in nonsense-mediated decay and several antiviral innate immune pathways (Figure S4B). Since human islets are constituted by several cell types, we asked whether the differentially expressed genes in human islets overlapped substantially with EndoC-βH1 cells. Intersection of the differentially expressed genes in human islets and EndoC-βH1 cells revealed 1577 and 1547 commonly upregulated and downregulated genes respectively (Figure S4C). Pathway analyses on the list resulting from the intersected upregulated and downregulated gene sets in human islets and EndoC-βH1 cells (FDR<0.05), revealed enrichment in pathways associated with non-sense mediated decay and the antiviral innate immune response (Figure S4D). The gene expression similarities between human islets (Figure S4E) and EndoC-βH1 (Figure S4F) were remarkable and showed that EndoC-βH1 cells recapitulate human islet transcriptomic remodeling induced by IL-1 β and IFN-α. Indeed, virtually all innate immune sensor genes, were consistently upregulated in both experiments (Figures S4E and S4F). Importantly, numerous innate immune genes commonly upregulated in human islets and EndoC-βH1 cells treated with IL-1β and IFN-α, overlapped substantially with those from the human insulitic islets from T1D onset patients, and notably included upregulation of *OAS1, OAS2,* and *OAS3* genes (Pedersen *et al.,* 2021).

Next, we explored the m^6^A regulation of the innate immune response by analyzing the m^6^A methylome of IL-1 β and IFN-α-treated EndoC-βH1 cells. We then confirmed that the m^6^A peaks are enriched at the start and stop codons (Figure S4G) and characterized by the canonical GGACU motif (Figure S4H). Analysis of m^6^A-sequencing revealed 301 differently methylated sites in 182 genes (FDR<0.05) and a higher number of sites with increased levels of m^6^A methylation in IL-1 β and IFN-α-treated compared to PBS-treated samples (Figure S4I).

To further explore and confirm the m^6^A regulation of the OAS innate immune response in β-cells, we intersected the commonly expressed and m^6^A methylated genes of human islets or EndoC-βH1 cells treated with IL-1 β and IFN-α versus PBS treatment (Figure 4D). This analysis identified 36 commonly expressed and m^6^A-regulated genes mainly involved in the innate immune pathway including the OAS antiviral response genes (Figure 4E). Protein-protein interaction analyses on the common m^6^A-regulated and expressed genes revealed a close relationship among OAS1, OAS2, and OAS3 proteins with other innate immune mediators (Figure 4F). Importantly, we were able to validate the m^6^A hypermethylation of OAS1 and OAS2 in EndoC-βH1 cells challenged with IL-1β and IFN-α compared to PBS (Figure 4G). Together, these results point to the m^6^A regulation of the OAS innate immune gene expression response in β-cells at T1D onset.

### m^6^A mRNA methylation promotes the decay of OAS genes in human β-cells to regulate the innate immune response at T1D onset

The observation that cytokine stimulation of β-cells upregulates the mRNA and the m^6^A levels of OAS genes prompted us to hypothesize that m^6^A directly regulates the upregulation of OAS protein levels. To address this hypothesis, we challenged EndoC-βH1 cells harboring METTL3 knock-down (KD) or scramble with IL-1 β and IFN-α or PBS for 16h (Figure 4H). Consistent with an earlier report that OAS protein levels are highly induced by interferon signaling and nearly absent at basal states (Marie et al., 1990) we observed that OAS1, OAS2, and OAS3 were upregulated in scramble cells stimulated with cytokines compared to PBS (Figure 4H). Furthermore, METTL3 silencing led to a further increase in OAS proteins and downstream innate immune mediators such as RNASE L in response to cytokines (Figure 4H and I). To validate this finding in primary cells, we dispersed human islets, transfected them with scramble or siMETTL3, and treated the re-aggregated pseudoislets with IL-1 β and IFN-α or PBS for 16h (Figure 4J). Pseudoislets with METTL3 silencing exhibited a greater upregulation of OAS proteins (Figure 4J and 4K) suggesting that low METTL3 levels allow an exacerbated OAS response in human β-cells in response to IL-1 β and IFN-α.

We hypothesized that METTL3 controls the innate immune response in β-cells by accelerating the mRNA decay of OAS genes, and that METTL3 downregulation leads to a sustained OAS activation, unresolved innate immune response, and T1D. To test this, we challenged EndoC-βH1 cells harboring METTL3 KD or scramble with actinomycin D (a transcription inhibitor) or DMSO in the presence of IL-1β and IFN-α, and harvested cells at various time points for mRNA analyses (Figure 4L). We observed that METTL3 silencing increases the mRNA stability of OAS genes compared to scramble cells in response to cytokines (Figure 4L). To validate these findings, we asked whether the converse, i.e. upregulating METTL3 leads to accelerated decay of OAS genes. Indeed, overexpression of METTL3 in EndoC-βH1 cells followed by a time-course treatment with actinomycin D in IL-1 β and IFN-α-treated cells led to an accelerated decay of OAS genes in response to cytokines compared to control (FLAG overexpressing) cells (Figure 4M).

To validate the m^6^A regulation of *OAS* expression and identify the putative m^6^A reader proteins controlling OAS mRNA decay at T1D onset, we silenced YTHDF1, YTHDF2, or YTHDF3 independently in EndoC-βH1 cells treated with IL-1 β and IFN-α and analyzed the decay dynamics of OAS1, OAS2, and OAS3 mRNAs upon actinomycin D treatment (Figure 4N). YTHDF1 or YTHDF3 downregulation increased *OAS* mRNA stability, suggesting that m^6^A hypermethylation of *OAS* transcripts accelerates the decay of their mRNA mediated by YTHDF1 and YTHDF3 (Figure 4N). In summary, these data reveal that METTL3 upregulation and consequent m^6^A hypermethylation of *OAS1, OAS2,* and *OAS3* control the innate immune response in human β-cells by promoting its mRNA decay via YTHDF1 and YTHDF3 (Figure 4O).

### m^6^A landscape of established T1D is enriched in β-cell identity and function genes

To explore the m^6^A landscape of established T1D, we performed RNA-seq and m^6^A-seq in human islets from patients with established T1D or non-diabetic controls (Figure 5A) (patient information in Supplementary Table 1). The transcriptome (Figure 5B) and the m^6^A methylome (Figure 5C) of established T1D versus non-diabetic control islets exhibited clear segregation in the principal component analysis (PCA). T1D islets presented downregulation of 5441 genes and an upregulation of 4913 genes compared to controls (FDR<0.05) (Figure 5D). Pathway enrichment analyses of the combined upregulated and downregulated genes (FDR<0.05) (Figure 5D) revealed pathways involved in translation initiation, nonsense-mediated decay, insulin secretion, and maturity-onset of diabetes of the young (MODY)(Figure 5E).

**Figure 5:**
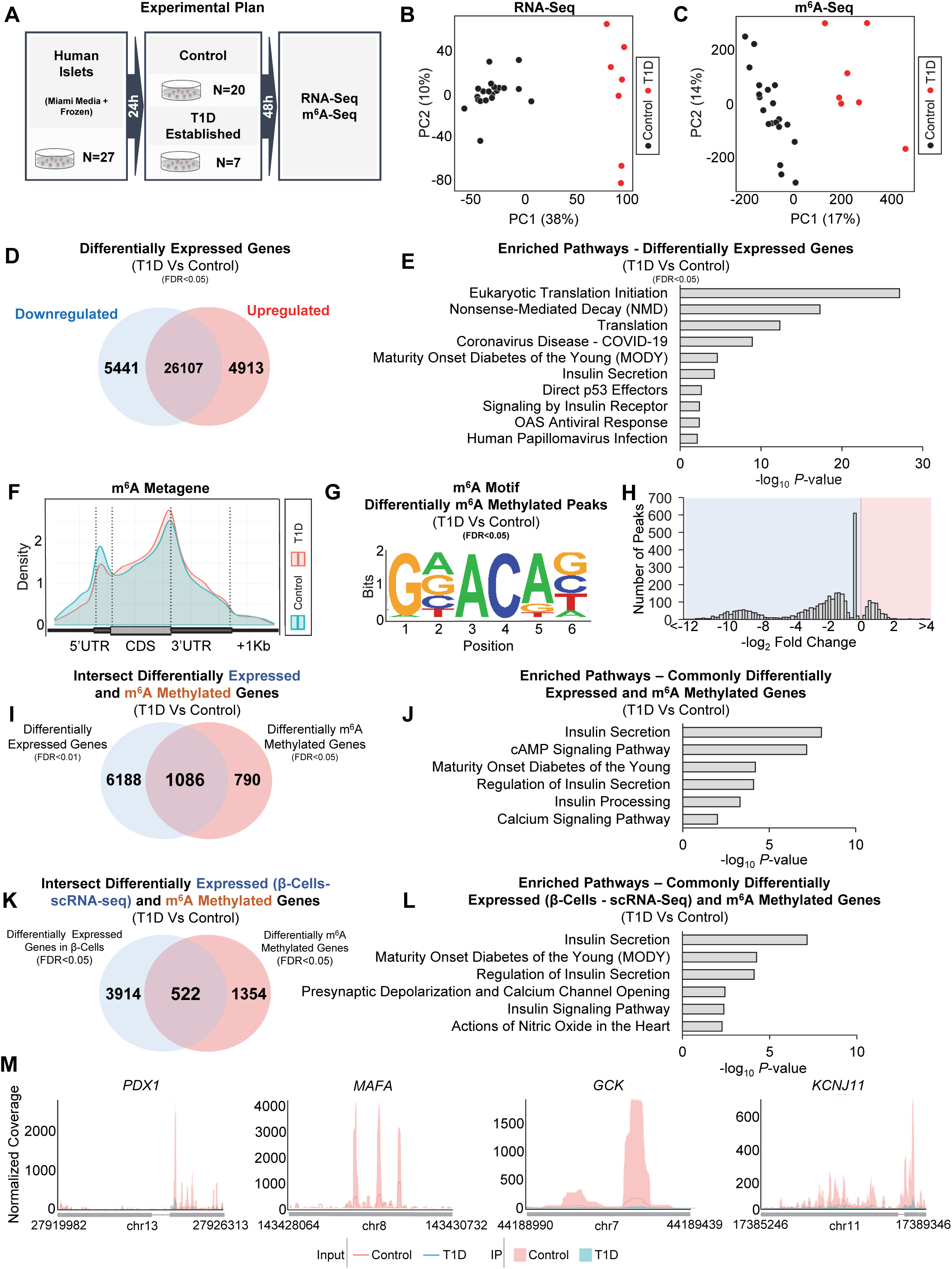
m^6^A landscape of established T1D is enriched in β-cell identity and function genes. **(A)** Summary scheme of the experimental plan. **(B)** PCA plot of RNA-sequencing in human islets from Control (black dots) or established T1D (red dots). **(C)** PCA plot of m^6^A-in human islets from Control (black dots) or established T1D (red dots). **(D)** Venn diagram representation of the upregulated (red), downregulated (blue), and unchanged genes (black) in human islets from Control (black dots) or T1D (red dots). **(E)** Pathway enrichment analyses of upregulated and downregulated genes in human islets from T1D compared to Controls. P-values were calculated according to the hypergeometric test based on the number of physical entities present in both the predefined set and user-specified list of physical entities. **(F)** Metagene of m^6^A enriched peaks in Control (blue) and T1D (red) human islets. **(G)** Enrichment for known m^6^A consensus motif RRACH. **(H)** Histogram showing the distribution of differential m^6^A loci log_2_ fold changes from T1D versus Control human islets. **(I)** Intersection of differentially expressed and m^6^A methylated genes in T1D compared to Control human islets. **(J)** Pathway enrichment analyses of intersected genes in (I). **(K)** Intersection of differentially expressed genes in T1D β-cells compared to Control from a published scRNA-seq dataset (Russell *et al.,* 2019) and our m^6^A dataset comparing the differentially m^6^A methylated genes in established T1D compared to Control human islets. **(L)** Pathway enrichment analyses of intersected genes in (K). **(M)** Coverage plots of m^6^A peaks in β-cell identity genes in human islets from established T1D compared to Controls. Plotted coverages are the median of the n replicates presented. Human islets: Controls n=20 and T1D n=7 biologically independent samples. EndoC-βH1 cells: n=6 biologically independent samples. Statistical analyses were performed using the Benjamini-Hochberg procedure.

m^6^A-seq analysis confirmed the enrichment of m^6^A peaks at the start and stop codons (Figure 5F) and were characterized by the canonical GGACU motif (Figure 5G). Our study identified 3485 differently methylated sites in 1876 genes (FDR<0.05) and a higher number of sites presenting m^6^A hypomethylation in established T1D compared to controls (Figure 5H) consistent with our findings of downregulation of METTL3 with T1D progression (Figure 1E and F).

To dissect the m^6^A regulation of gene expression in established T1D, we intersected and performed pathway enrichment analysis on the differentially expressed and m^6^A methylated genes in human islets from patients with established T1D compared to controls (Figure 5I). This analysis identified enrichment for pathways associated with β-cell function and identity such as insulin secretion, cAMP signaling, MODY, insulin processing, and calcium signaling (Figure 5J). Since human islets are constituted by several cell types and the number of insulin-positive β-cells is drastically decreased in established T1D, we intersected the differentially m^6^A decorated genes performed in whole islets of established T1D, with the β-cell transcriptome from a single-cell RNA seq dataset performed in islets from an independent cohort of established T1D patients and non-diabetic controls (GSE121863) (Figure 5K). Pathway analyses on the intersected differentially expressed and m^6^A methylated genes revealed enrichment for insulin secretion, and MODY (Figure 5L). Among the m^6^A hypomethylated genes in established T1D were master regulators of β-cell function and identity including *PDX1, MAFA, GCK,* and *KCNJ11* (Figure 5M). Together, these results point to distinct m^6^A landscapes at T1D onset and established T1D in human β-cells. Our results point to the m^6^A regulation of β-cell function and identity genes in established T1D.

### Sustained over-expression of Mettl3 in β-cells delays diabetes progression in the NOD mouse model of T1D

Several lines of evidence implicate the involvement of the innate immune system in β-cell destruction and the development of T1D (Tai et al., 2016). For example, altered type I interferon-induced gene signature is a common feature of T1D onset in both humans and NOD mice (Akazawa et al., 2021).

We hypothesized that an intervention that promotes a sustained upregulation of Mettl3 in the NOD mouse β-cells after T1D onset, would lead to a faster turnover and/or decrease in expression of Oas and protect β-cells to delay T1D. To test this, we designed two different adeno-associated virus serotype 8 (AAV8) driving eGFP or Mettl3 under the control of the rat insulin promoter II (RIP2) that has been previously used to achieve β-cell specificity (Dor et al., 2004) (Figure 6A). We chose to infuse PBS or AAV8 into 4-week-old animals (Figure 6B) because our data showed that Mettl3 levels start to decline after this age in NOD females (Figure 1A and 1H). We confirmed Mettl3 protein overexpression at 12 weeks of age (Figure 6C) and followed these mice for up to 25 weeks of age similar to previous studies in the NOD mouse model (Kulkarni et al., 2021; Nelson et al., 2020; Rui et al., 2021). Body weight trajectories didn’t change until 17 weeks of age when mice that received AAV8 driving eGFP overexpression (AAV8-eGFP) or PBS started to lose weight compared to mice that received AAV8 driving Mettl3 overexpression (AAV8-Mettl3) (Figure 6D). Random-fed blood glucose levels increased with age in AAV8-eGFP and PBS groups, in contrast to the AAV8-Mettl3 group which presented an improved glycemic profile (Figure 6E). Thus, Mettl3 overexpression delayed T1D compared to AAV8-eGFP or PBS (Figure 6F).

**Figure 6:**
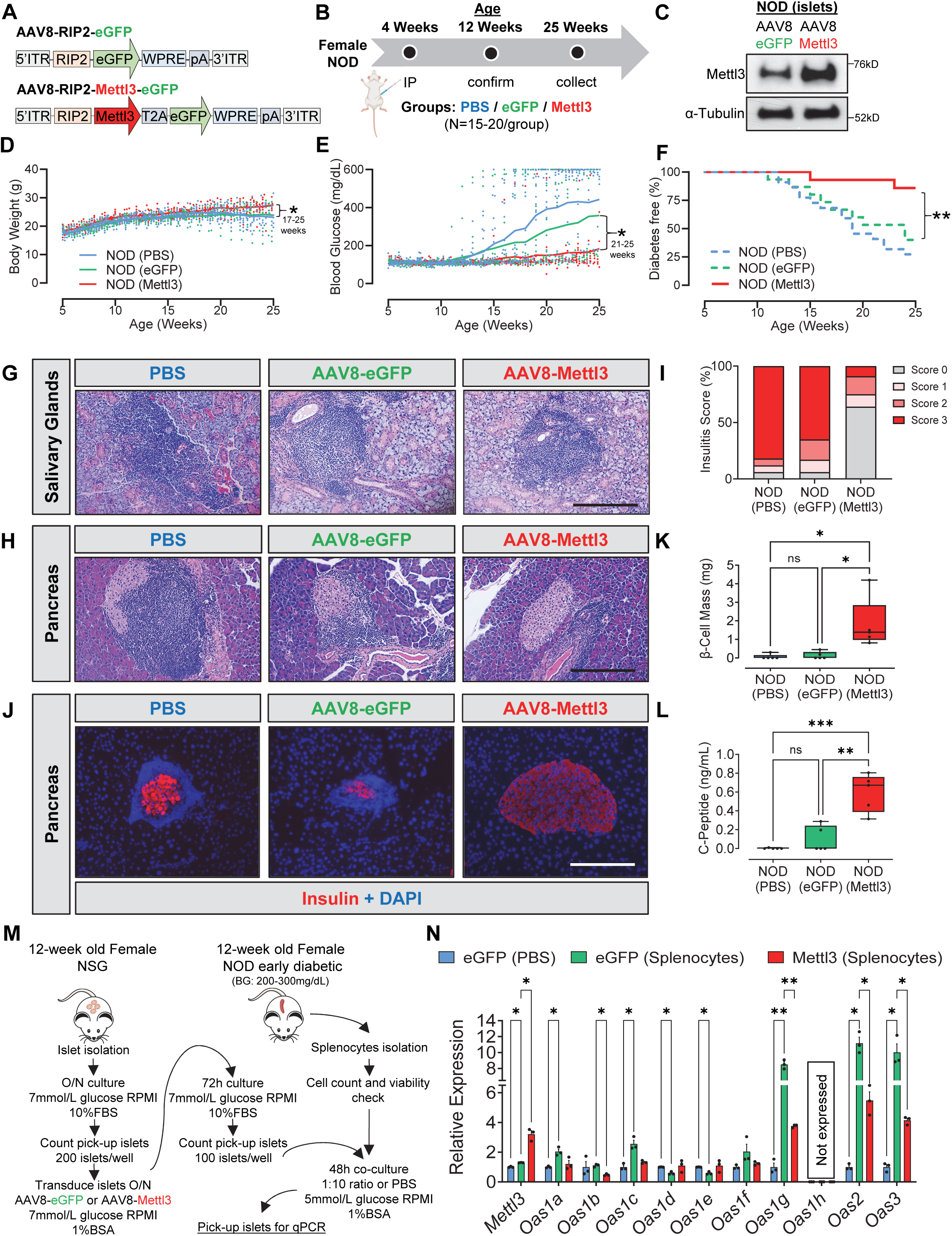
*In vivo* AAV8-mediated overexpression of Mettl3 in NOD β-Cells delays Type 1 diabetes progression. **(A)** Schematic diagram showing the construction of AAV8 driving eGFP or Mettl3 under the control of a rat insulin promoter II. **(B)** Scheme of experimental approach depicting NOD mice receiving PBS (blue), AAV8 overexpressing eGFP (green), or AAV8 overexpressing Mettl3 (red). N=20 in NOD (PBS), and n=15 in NOD (eGFP) or NOD (Mettl3). **(C)** Western-blot validation of Mettl3 overexpression in isolated NOD female islets after 8 weeks of *in vivo* transduction. **(D)** Body weight trajectories of NOD (PBS) (blue dots/line), NOD (eGFP) (green dots/line), and NOD (Mettl3) (red dots/line). **(E)** Blood glucose trajectories of NOD (PBS) (blue dots/line), NOD (eGFP) (green dots/line), and NOD (Mettl3) (red dots/line). **(F)** Percentage diabetes-free NOD (PBS), NOD (eGFP), and NOD (Mettl3) at the end of the 25 weeks of age. **(G)** Representative hematoxylin and eosin (H&E) staining showing immune cell infiltration in salivary glands in NOD (PBS), NOD (eGFP), or NOD (Mettl3) (scale bar, 200 µm). **(H)** Representative H&E staining showing immune cell infiltration in pancreatic islets in NOD (PBS), NOD (eGFP), or NOD (Mettl3) (scale bar, 200 µm). **(I)** Quantification of insulitis score of pancreatic sections from H (n=5/group). **(J)** Representative immunofluorescence images showing insulin (red) and DAPI (blue) in pancreatic sections from NOD (PBS), NOD (eGFP), or NOD (Mettl3) (scale bar, 200 µm). **(K)** β-cell mass estimations of NOD (PBS), NOD (eGFP), and NOD (Mettl3) at 25 weeks of age (n=5/group). **(L)** Serum C-peptide levels in NOD (PBS), NOD (eGFP), and NOD (Mettl3) at 25 weeks of age (n=5/group). **(M)** Schematic representation of the co-culture experimental plan. **(N)** qRT-PCR analyses of Oas genes in NOD *scid* gamma (NSG) islets transduced with eGFP and co-cultured with PBS (blue bars) or NOD diabetogenic splenocytes (green bars), or NSG islets transduced with AAV8 overexpressing Mettl3 and co-cultured with NOD diabetogenic splenocytes (red bars) (n=3/group; islets from 3 pools of 5 mice each pool). All samples in each panel are biologically independent. Data were expressed as means ± SEM. \**P*<0.01, \*\**P*<0.01, \*\*\**P*<0.001. Statistical analysis was performed by multiple paired t-tests in D and E. Log-rank (Mantel-cox) test in F. Two-Way ANOVA with Holm-Sidak’s multiple comparisons test in K, L, and N.

NOD mice present immune cell infiltration in salivary glands as well as pancreatic islets during progression of T1D (Robinson et al., 1996). Assessment of immune cell infiltration in these two tissues helps evaluating if a potential therapeutic approach induces β-cell specific or systemic immune protection (Dirice et al., 2019). While all groups presented salivary gland infiltration (Figure 6G), only AAV8-Mettl3 treated mice showed decreased immune infiltration in pancreatic islets (Figure 6H) and consequent decreased insulitis scores compared to AAV8-eGFP or PBS-treated groups (Figure 6I). Consistently, AAV8-Mettl3 showed increased β-cell mass (Figure 5J and K), and serum C-peptide levels compared to AAV8-eGFP or PBS (Figure 6L). Thus, sustained upregulation of Mettl3 protects β-cells and delays T1D in the NOD mouse and points to the enzyme as an attractive therapeutic target.

### Mettl3 overexpression hampers Oas immune response in NOD islets

Our studies in human islets and β-cells revealed that METTL3 controls OAS response to IL-1 β and IFN-α by regulating mRNA decay via YTHDF1 and YTHDF3. To test the hypothesis that Mettl3 overexpression in mouse β-cells limits Oas upregulation in response to a T1D immune insult we employed co-culture experiments (Figure 6M). For this, we first transduced islets from 12-week-old female immunodeficient NOD SCID gamma (NSG) mice with AAV8-eGFP or AAV8-Mettl3 and then co-cultured them with PBS or diabetogenic splenocytes from 12-week old NOD females that recently developed diabetes (Figure 6M). Co-culture of islets from NSG mice that were transduced with AAV8-eGFP with diabetogenic splenocytes increased *Mettl3* compared to AAV8-eGFP treated with PBS (Figure 6N). These data suggest that the cytokines secreted by diabetogenic immune cells initially upregulate Mettl3 in mouse islets. In addition, AAV8-Mettl3-islets presented a greater Mettl3 upregulation compared to AAV-eGFP-islets when challenged with splenocytes (Figure 6N) demonstrating that *Mettl3* overexpression *in vitro* was successful. *Oas1* exhibits 8 paralog genes which have been described to differ in their antiviral activity (Elkhateeb et al., 2016). Co-culture of diabetogenic splenocytes with islets transduced with AAV8-eGFP induced the upregulation of *Oasla, Oaslc, Oaslg, Oas2,* and *Oas3* compared to islets transduced with AAV8-eGFP that were challenged with PBS (Figure 6N). On the other hand, overexpression of Mettl3 (AAV8-Mettl3) in β-cells prior to co-culture with diabetogenic splenocytes blunted the upregulation of Oas genes compared to β-cells transduced with AAV8-eGFP and co-cultured with diabetogenic splenocytes (Figure 6N). These data provide strong evidence for the existence of a conserved METTL3 regulation of the OAS innate immune response in mouse and human β-cells.

### ROS-induced oxidative stress and endoplasmic reticulum (ER) stress constrain METTL3 upregulation in human islets in response to IL-1 β and IFN-α

To examine the mechanisms involved in the dynamic regulation of METTL3 in T1D we considered recent reports describing the involvement of mitochondrial dysfunction, reactive oxygen species (ROS), and ER stress as contributors to the development of disease (Chen et al., 2018a; Kim et al., 2021; Tersey et al., 2012).

First, we challenged human islets with PBS, a ROS inducer (hydrogen peroxide-H_2_O_2_) or H_2_O_2_ in the presence of a ROS scavenger (N-acetyl cysteine - NAC) (Figure S5A). Human islets treated with H_2_O_2_ presented downregulation of thioredoxin-1 (TXN), which is degraded by H_2_O_2_ as previously reported (Haendeler et al., 2005) (Figure 7A). METTL3 protein levels were significantly downregulated by H_2_O_2_ and this downregulation was rescued by co-treatment with NAC (Figure 7A and 7B). Together these data suggest that pathological levels of ROS, such as those present in established T1D, can downregulate METTL3. Next, to examine the impact of ER stress on METTL3 regulation in the context of T1D, we treated human islets with DMSO, or DMSO followed by IL-1 β and IFN-α. Alternatively, we incubated human islets with thapsigargin (an ER stress inducer) prior to treatment with IL-1 β and IFN-α (Figure S5B). Consistent with our previous results (Figure 2D), cytokine treatment led to the upregulation of all three components of the m^6^A writer complex, METTL3, METTL14, and WTAP (Figure 7C and 7D). However, induction of ER stress, by prior incubation with thapsigargin, blunted METTL3 upregulation and also blocked the increase in METTL14 and WTAP in response to IL-1 β and IFN-α treatment (Figure 7C and 7D). These results suggest that the accumulation of ROS and the presence of ER-stress dampens METTL3 upregulation in response to cytokines.

**Figure 7:**
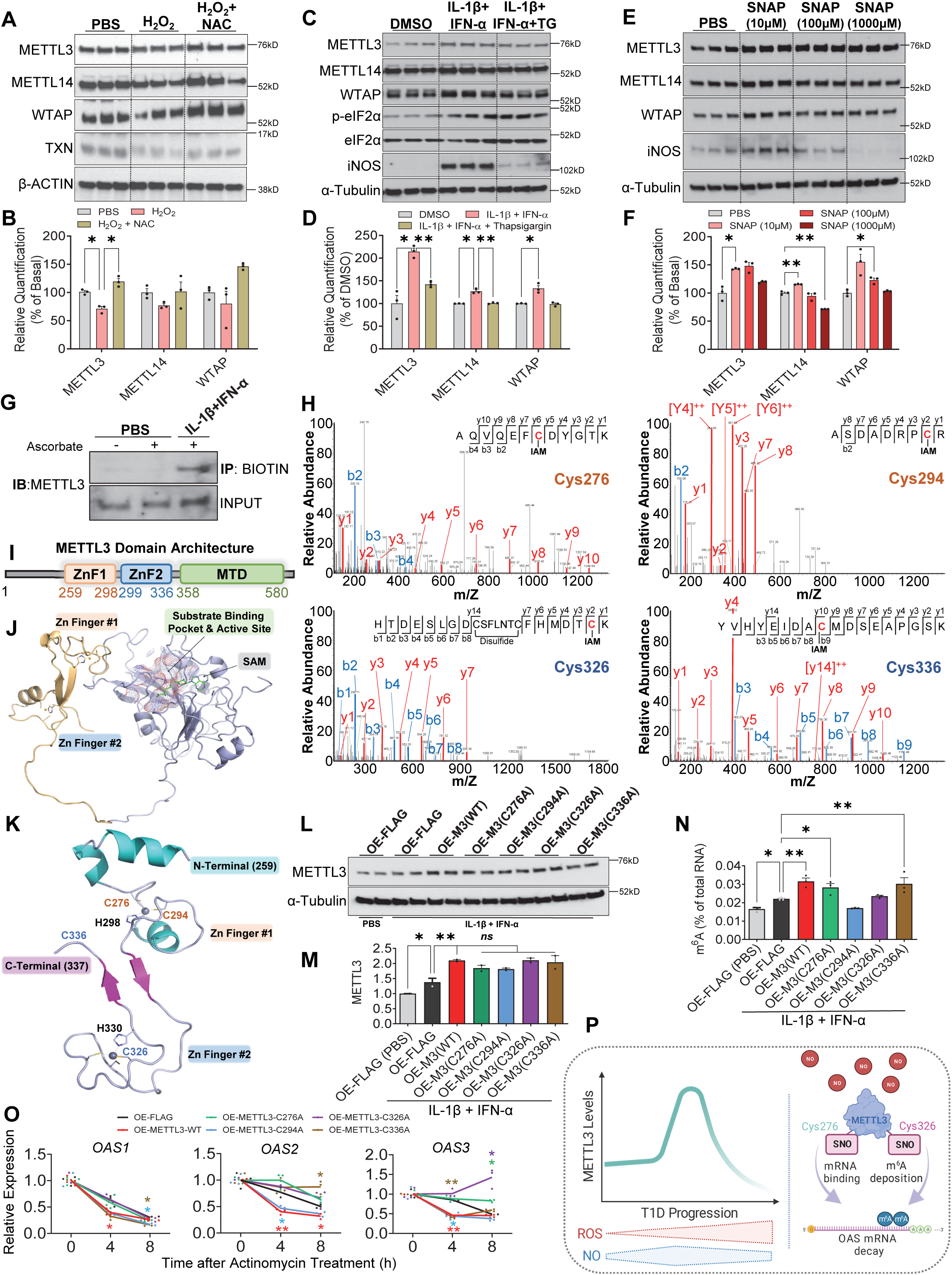
OAS mRNA stability is regulated by the S-nitrosylation of METTL3 in human β-cells. **(A)** Western-blot analyses of indicated proteins in human islets cells treated with H_2_O_2_ or H_2_O_2_ plus N-acetyl cysteine (NAC) for 24h (n=3/group). **(B)** Protein quantification of indicated proteins related to (A). **(C)** Western-blot analyses of indicated proteins in human islets treated with DMSO, IL1-β plus IFN-α pre-treated with thapsigargin plus IL1-β and IFN-α for a total of 24h (n=3/group). **(D)** Protein quantification of indicated proteins related to (C). **(E)** Western-blot analyses of indicated proteins in human islets treated with PBS or represented doses of S-nitroso-N-acetyl-DL-penicillamine (SNAP) for 24h (n=3/group). **(F)** Protein quantification of indicated proteins related to (E). **(G)** Western-blot analyses of biotin-switch assay on METTL3 in EndoC-βH1 cells treated with PBS or IL1-β plus IFN-α (n=3 experiments). **(H)** MS/MS spectra of S-nitrosylated METTL3 peptides and the identification of cysteines C276, C294, C326, and C336 (n=2 replicates). **(I)** METTL3 protein domains representing zinc finger domains (ZnF) and methyltransferase domain (MTD). **(J)** Structure of human METTL3 depicting two zinc finger domains and MTD. **(K)** Structure of METTL3 zinc finger domains depicting the identified cysteines sensitive to S-nitrosylation. **(L)** Western-blot analyses of METTL3 in EndoC-βH1 cells (n=2/group). **(M)** Protein quantification of METTL3 related to (L). **(N)** m^6^A levels measured by a colorimetric ELISA kit of total RNA isolated from EndoC-βH1 cells overexpressing the represented plasmids and treated with PBS or IL-1 β plus IFN-α for 48h (n=3/group). **(O)** qRT-PCR analyses of OAS genes after IL-1 β plus IFN-α stimulation in EndoC-βH1overexpressing the represented plasmids after a time-course treatment with ActD (n=3/group). **(P)** Model depicting the role of S-nitrosylation in controlling METTL3 function and OAS mRNA decay. All samples in each panel are biologically independent. Data were expressed as means ± SEM. Statistical analysis was performed by Two-Way ANOVA with Fisher’s LSD test.

While these data provide one explanation for the downregulation of METTL3 during progression of T1D, we next considered the mechanism(s) for the converse, i.e. METTL3 upregulation during the early stages of T1D. In this context, it is notable that pro-inflammatory cytokines induce nitric oxide synthesis in β-cells and while physiological levels of NO are protective, the concomitant persistent accumulation of ROS and nitric oxide to pathological levels may trigger β-cell apoptosis (Chen *et al.,* 2018a). Our data demonstrated that while human islets treated with IL-1 β and IFN-α exhibited robust upregulation of inducible nitric oxide synthase (iNOS) (Figure 7C and 7D), pre-treatment with thapsigargin blocked the response similar to non-ER-stressed islets (DMSO-treated) (Figure 7C and 7D).

To directly examine the involvement of NO in mediating METTL3 upregulation at T1D onset we challenged human islets with different concentrations of a NO donor (S-Nitroso-N-acetyl-DL-penicillamine-“SNAP”) (Figure S5C). Low levels of SNAP increased METTL3 and iNOS protein abundance, while this response was lost in islets treated with high doses of SNAP (Figure 7E and 7F). Altogether, these data support the concept that at early stages of T1D, an initial increase in NO induces upregulation of METTL3. However, as the disease progresses, the accumulation of NO and ROS in β-cells coupled with exacerbation of ER stress downregulates METTL3 and activates a persistent OAS response and consequent β-cell death.

### Regulation of OAS mRNA decay in β-cells is dependent on S-nitrosylation of cysteine residues (C276 and C326) in the zinc finger domain of METTL3

The redox modification of a cysteine (Cys) thiol by NO forms an S-nitrosothiol. Growing evidence points to S-nitrosylation (SNO) as a valuable therapeutic target to improve β-cell function (Zhou et al., 2022). To further explore the mechanism(s) involved in the regulation of METTL3 by NO, first, we employed a biotin-switch assay to confirm that METTL3 SNO levels were increased by IL-1 β and IFN-α in EndoC-βH1 cells (Figure 7G). Next, to identify the covalently modified Cys residues which were susceptible to SNO, we optimized a protocol of liquid chromatography with tandem mass spectrometry (LC-MS/MS) on human recombinant METTL3 protein treated with SNAP (a NO donor) (Figure S5D). These analyses identified the SNO of four evolutionary conserved METTL3 Cys residues (Cys276, Cys294, Cys326, and Cys336) (Figure 6H). The domain architecture of human METTTL3 is characterized by the existence of two zinc finger domains (CCCH) and a methyltransferase domain (MTD) (Wang *et al.,* 2016) (Figure 7I and 7J). Interestingly, all METTL3 Cys residues sensitive to SNO were located in the zinc fingers (Figure 7K).

CCCH-type zinc finger domains are known to be involved in RNA binding and immune regulation (Fu and Blackshear, 2017). Furthermore, SNO of zinc fingers may lead to reversible disruption of zinc finger structures and modulate their DNA and RNA binding capacities (Garban et al., 2005; Kroncke, 2001). We hypothesized that SNO of the identified cysteine residues were fundamental for the METTL3 regulation of OAS mRNA stability. To test this, we performed site-directed mutagenesis by substituting each of the four candidate cysteines with alanines since the latter is physicochemically innocuous and does not react with NO. Next, we overexpressed either wild-type (WT) METTL3 or METTL3 bearing mutant Cys constructs in EndoC-βH1 cells followed by treatment with IL-1 β and IFN-α or PBS for 24h. Cells overexpressing a FLAG empty plasmid treated with cytokines presented upregulation of METTL3 compared to PBS (Figure 7L and 7M). Overexpression of WT METTL3 in EndoC-βH1 cells was successful (Figure 7L and 7M). Importantly, all mutants upregulated METTL3 similarly to WT METTL3 compared to a FLAG empty plasmid (Figure 7L and 7M). However, while overexpression of WT METTL3 in EndoC-βH1 cells increased m^6^A levels in total RNA, this was blocked by mutating Cys294 and Cys326 (Figure 7N). This data shows that SNO at METTL3 Cys294 and Cys326 is important for m^6^A deposition.

Finally, to test which of the four cysteines directly influences METTL3 control of the decay process of OAS mRNA in β-cells we overexpressed FLAG, WT METTL3, or Cys mutant constructs independently in EndoC-βH1 cells and collected RNA at different time points after actinomycin D treatment in the presence of IL-1 β + IFN-α. Consistent with our previous data, METTL3 overexpression accelerated the mRNA decay of *OAS1, OAS2,* and *OAS3* compared to a FLAG control plasmid (Figure 7O). Overall, mutant C294A overexpression cells behaved similarly to WT METTL3 (Figure 7O), suggesting that SNO at this Cys residue does not control the mRNA decay of *OAS* despite impacting global m^6^A deposition in total RNA. Overexpression of C336A accelerated the mRNA decay of *OAS1* and *OAS3* similarly to WT METTL3. However, SNO at this specific Cys was necessary for the mRNA decay of *OAS2* (Figure 7O). On the other hand, overexpression of mutants C276A and C336A behaved similarly to FLAG overexpressing cells and did not accelerate OAS mRNA decay as seen in the WT METTL3 overexpressing cells (Figure 7O). Together, these data suggest that SNO regulates METTL3 and OAS mRNA decay by two different mechanisms. While SNO at Cys 326 impacts METTL3 global m^6^A deposition capacity, Cys 276 is likely important for the OAS RNA binding to METTL3 and does not disturb METTL3 m^6^A deposition in total RNA (Figure 7P).

## DISCUSSION

There is growing evidence for a central role of the β-cell in triggering autoimmunity in T1D. These observations have been summarized recently (Mallone and Eizirik, 2020) and include: a) heterogeneity of insulitis observed in T1D pancreases; b) the identification of multiple T1D-associated GWAS variants expressed by β-cells, c) the presence of ER stress and metabolic abnormalities in β-cells that precede T1D seroconversion. While these reports collectively argue for a compromised dialogue between β-cells and the immune system that might trigger T1D pathogenesis, the precise mechanism(s) directing the mismatch in this communication remain unclear. Here, we report novel findings that the m^6^A writer, METTL3, is dynamically regulated in β-cells and that m^6^A mRNA methylation is important in controlling the antiviral innate immune response at the onset of T1D. Moreover, we note SNO as a novel post-translational regulatory mechanism for METTL3 function and OAS mRNA decay as a component of the pathogenic process.

To directly evaluate the biological relevance of the METTL3 upregulation in β-cells at the onset of T1D and considering the inaccessibility of human islets from living patients with T1D, we designed an *in vitro* approach to recapitulate the phenotype that mimics early onset of the disease. Specifically, we utilized cytokines, such as interleukin-1 β (IL-1 β) and interferon α (IFN-α), that have long been noted to mediate the inflammation associated with the accumulation of immune cells during the early stages of T1D (Eizirik *et al.,* 2009; Todd, 2010). Indeed, these cytokines have been used extensively to recapitulate several pathophysiological aspects of T1D (Colli *et al.,* 2018; Colli et al., 2020; Eizirik *et al.,* 2012; Ramos-Rodriguez *et al.,* 2019). Consistently, we observed that IL-1β and IFN-α treatment increased m^6^A levels and METTL3 in human β-cells. This allowed us to map the m^6^A landscape of T1D onset in human β-cells by performing m^6^A-seq and RNA-seq in 15 independent human islet preparations and six EndoC-βH1 cell batches treated with IL-1 β and IFN-α or PBS.

Cytokine treatment induced profound transcriptomic remodeling in human islets and EndoC-βH1 cells and mostly impacted innate immune networks, including the upregulation of the antiviral OAS innate immune pathway. Importantly, these changes were consistent with the transcriptional remodeling described in laser-captured micro-dissected (LCM) insulitic islets from patients recently diagnosed with T1D (Lundberg *et al.,* 2016; Pedersen *et al.,* 2021). These data argue that cytokine treatment recapitulated the transcriptomic signatures of human islets at T1D onset and identified OAS genes to be upregulated at early stages of disease.

m^6^A-seq analyses revealed hypermethylation of several innate immune sensors, including OAS genes in response to cytokine treatment in β-cells. Herein, we present several lines of evidence to validate the ability of METTL3 to control the mRNA decay of OAS genes via the m^6^A pathway. First, silencing *METTL3* in EndoC-βH1 cells or human pseudoislets increased OAS protein levels in response to cytokine treatment. Second, EndoC-βH1 cells deficient for METTL3, exhibited increased stability of OAS mRNA while conversely overexpression of METTL3 accelerated their mRNA decay. Third, we demonstrated that the mRNA decay of OAS genes in EndoC-βH1 cells is mediated by the m^6^A readers YTHDF1 and YTHDF3. Finally, we demonstrated that the β-cell overexpression of Mettl3 in mice *ex vivo,* dampened Oas upregulation in response to diabetogenic splenocytes.

Innate immune sensors are in constant contact with endogenous and exogenous nucleic acids. One proposed evolutionary consequence of repeated viral infections in humans is the development of exuberant and sensitive antiviral innate immune responses to counterbalance the ability of viruses to mutate (Crowl *et al.,* 2017). Thus, it is not surprising that several autoimmune diseases have associated dysregulation of innate immune mediators (Carry *et al.,* 2022; Lang *et al.,* 2007; Tai *et al.,* 2016). For example, OAS proteins have been identified as enzymes that sense non-self nucleic acids and capable of initiating antiviral pathways, and with respect to these studies such proteins have been genetically linked with T1D, with *OAS1* variants being reported to increase disease risk (Pedersen *et al.,* 2021). Indeed, OAS genes are highly upregulated in human islets specifically at T1D onset (Lundberg *et al.,* 2016; Pedersen *et al.,* 2021). Investigations performed in rodent β-cell lines argue for a cell-specific effect with a 6-fold upregulation of OAS compared to α-cells in response to IFN-α or polyinosinic:polycytidylic acid [poly(I:C)] (dsRNA mimetic) (Dan *et al.,* 2012; Li *et al.,* 2009b). Consistently, constitutive expression of OAS in β-cells leads to apoptosis and inhibition of proliferation (Dan *et al.,* 2012; Li *et al.,* 2009b).

The repeated interactions with viruses may have resulted in the development of safeguard mechanisms in host cells to integrate activation thresholds for innate immune sensors to eliminate and/or prevent immune-stimulatory endogenous nucleic acids coupled with the temporal control of the response to exogenous nucleic acids. The virtual absence of reports on such mechanisms in β-cells identifies m^6^A as a new safeguard mechanism to control the innate OAS antiviral immune response and establishes a novel link between innate immunity and T1D. Recent reports support this concept. For example, Gao and colleagues show that loss of Mettl3 in fetal livers promotes the formation of deleterious dsRNAs and activation of innate immune pathways in the absence of a viral infection (Gao *et al.,* 2020). On the other hand, deletion of METTL3 (Winkler *et al.,* 2019) or METTL14 (Rubio et al., 2018) in fibroblasts increases the mRNA stability of interferon β upon viral infection. Finally, the virus strain can influence the ability of host cell m^6^A levels to modulate viral proliferation potential and recognition by innate immune sensors (Gokhale et al., 2020; Lu et al., 2020). Given the phenotypic heterogeneity of T1D, it is likely that these mechanisms variably impact β-cells in different patients. Further studies are needed to elucidate if m^6^A levels impact the β-cell susceptibility to viral infection and/or if downregulation of m^6^A levels in β-cells can trigger the formation of aberrant endogenous immune-stimulatory dsRNAs.

The m^6^A landscape of β-cells at T1D onset seemed to be distinct from established T1D. Intersection of differentially expressed genes from bulk or β-cell single-cell RNA seq with differentially m^6^A methylated genes in whole islets from established T1D revealed enrichment for β-cell identity and function pathways. This argues for distinct features between the transcriptome and m^6^A landscape of β-cells at T1D onset, such as observed in the human islets and EndoC-βH1 cells treated with cytokines, compared to β-cells from patients with established T1D. This suggests that as T1D progresses, m^6^A decorates and controls genes important for β-cell function and identity. Indeed, our previous work also identified m^6^A hypomethylation of several β-cell identity and function genes in human T2D islets compared to non-diabetic islets. Furthermore, Mettl14 deficient β-cells, which displayed decreased m^6^A levels, presented downregulation of several β-cell identity genes (De Jesus *et al.,* 2019). Overall, these results point to m^6^A as a mechanism involved in the loss of β-cell identity and function in the later stages of T1D.

To begin to dissect the mechanism(s) involved in the dynamic regulation of METTL3 during T1D progression, we considered recent findings that prompted the notion that the β-cell destruction is driven by their own high metabolic rate and low expression of anti-oxidant (e.g. SOD) and anti-apoptotic (e.g. BCL-2) proteins (Roep *et al.,* 2021). Indeed, increased accumulation of intracellular ROS has been reported in children prior to the development of T1D (Balzano-Nogueira et al., 2021). Furthermore, ER stress precedes T1D onset in NOD mice (Tersey *et al.,* 2012) and several ER-stress markers are known to be upregulated in human T1D β-cells (Eizirik et al., 2013). Our studies added important details to such notions in their revelation that increased levels of ROS and ER stress diminished METTL3 upregulation in human islets in response to cytokines, consistent with the hypothesis that β-cell redox state and ER stress impact METTL3 levels during T1D progression. This represents one possible explanation for the decrease in METTL3 levels during progression of the disease. On the other hand, while physiological levels of NO and cytokines, led to the upregulation of iNOS and METTL3, pathological levels of NO restored iNOS and METTL3 proteins to non-stimulated levels within 24h of treatment. The increase in METTL3 levels at the onset of the disease is likely due to redox sensitivity of METTL3 that responds to graded levels of ambient NO. Thus, during the initial stages of T1D, when an immune attack and cytokine release occurs, the rising levels of NO are still within the physiological range and promote upregulation of METTL3. However, as the disease progresses with gradual accumulation of NO and ROS, the heightened redox sensitivity of METTL3 leads to its downregulation.

The upregulation of OAS genes at the onset of T1D, despite an increase in METTL3, could be due to stressed β-cells failing to timely increase expression of the m^6^A writer to counterbalance the rapid and robust rise of OAS, which is in contrast to healthy cells. The fact that human islets pre-exposed to ER stress by thapsigargin treatment do not upregulate METTL3 similarly to control islets support this contention. Furthermore, in *in vitro* co-culture experiments where we boosted Mettl3 overexpression in β-cells, such efforts were adequate to limit the upregulation of Oas in islets from immunodeficient NSG mice in response to diabetogenic splenocytes from early diabetic age- and gender-matched NOD mice.

Finally, other mechanisms could also regulate the METTL3 m^6^A deposition independent of its total protein levels. For example, SUMOylation of METTL3 does not alter its stability, localization, or interaction with other m^6^A writers, but significantly represses its m^6^A methyltransferase activity (Du *et al.,* 2018). SNO has been reported to be essential for diverse aspects of β-cell function (Zhou *et al.,* 2022). We identified all four cysteines that are modified by SNO to be in the zinc finger domain of METTL3. This gains significance since SNO of zinc fingers has been reported to disrupt their structures and modulate enzymatic activity impacting RNA binding capacities of RNA binding proteins (RBPs) (Garban *et al.,* 2005; Kroncke, 2001). A recent study has also reported that the cysteines 294 and 326 in the zinc finger domains of METTL3 are essential for its enzymatic activity (Huang et al., 2019). We observed that the SNO of the METTL3 Cys276 and Cys326, is needed for the mRNA decay of OAS in response to cytokines. Interestingly, while mutation of Cys326 impacted m^6^A deposition, mutation of Cys276 did not alter m^6^A levels. This suggests that while SNO of Cys326 controls METTL3 enzymatic activity, SNO of Cys276 might regulate METTL3 structural binding to OAS mRNA. Overall, these results demonstrate that redox signaling controls METTL3 and OAS mRNA stability in β-cells at T1D onset.

In summary, we have uncovered evidence that m^6^A acts as a β-cell protective mechanism to control the OAS innate immune response at the onset of T1D in mice and humans. Our data also suggests that increased m^6^A promotes accelerated mRNA decay of OAS genes in β-cells. Importantly. we observed that SNO represents a previously unidentified mechanism with the capacity to modulate METTL3 protein function and potential mRNA binding affinity to OAS mRNA. Based on these results, we firmly believe that therapeutic targeting of METTL3 before seroconversion or at T1D onset has the potential to protect β-cells and to promote β-cell survival and function during disease progression.

## LIMITATIONS OF THE STUDY

In this study, we have shown that m^6^A is an adaptive β-cell protective mechanism that accelerates the mRNA decay of the 2’-5’-oligoadenylate synthetase (OAS) genes to control the antiviral innate immune response. We focused on mRNA decay since OAS mRNA levels were elevated in human islets and EndoC-βH1 cells treated with cytokines. Furthermore, this was consistent with the upregulation of *OAS* in human pancreas at T1D onset (Lundberg *et al.,* 2016; Pedersen *et al.,* 2021). Given that m^6^A regulates several aspects of the life of mRNA including RNA transcription rate and mRNA export from the nucleus to the cytoplasm, we cannot exclude that m^6^A might regulate OAS by additional mechanisms. Although the observations involving co-treatment of human islets and EndoC-βH1 cells with IL-1β and IFN-α were recapitulated to a vast extent by examining the β-cell transcriptomic landscape of human T1D onset, including the upregulation of several innate immune sensors including *OAS*, we also cannot exclude that other cytokine combination would reveal additional m^6^A methylated genes. Additionally, recent reports have pointed to the existence of cellular dysfunction at early stages of T1D impacting other pancreatic cell types, including α (Doliba et al., 2022) and acinar cells (Wright et al., 2020). Considering the widespread expression of m^6^A modulators in virtually all mammalian cells, it is plausible that m^6^A affects other pancreatic cell types during the onset of T1D. Taken together, we believe our novel investigations presented here provide strong rationale to pursue further studies to determine the impact of different cytokines in the m^6^A landscape of human β-cells and other pancreatic cell types. Our m^6^A-seq experiments in EndoC-βH1 cells detected a lower number of m^6^A peaks in comparison to human islets consistent with our previous mechanistic study (De Jesus *et al.,* 2019). We believe any such variance can be explained by EndoC-βH1 cells being biologically less differentiated and complex compared to human islets, and because we are employing a smaller number of replicates in comparison to human islets. Lastly, the fact that EndoC-βH1 cells are difficult to transfect cells and our previous work demonstrated that stable silencing of METTL3 leads to severe cell-cycle arrest (De Jesus *et al.,* 2019), prompted us to rely on transient KD approaches.

## DATA AND CODE AVAILABILITY

All data reported in this paper will be shared by the lead contact upon request. This paper does not report original code needed to reanalyze the data generated by this study. Any additional information required to reanalyze the data reported in this paper is available from the lead contact upon request.

## ACKNOWLEDGEMENTS

The authors thank Fatima Bosch PhD (Universitat Autònoma de Barcelona) and her lab members for discussions about AAV design and *in vivo* administration. We thank Anne Op de Beeck PhD (ULB Center for Diabetes Research, Université Libre de Bruxelles) for helpful discussions related to viral infections. We thank Ercument Dirice PhD (New York Medical College) for discussions related to co-culture of islets and splenocytes. We thank Britta Kunkemoeller PhD (Brigham and Women’s Hospital) for assistance with FACS analysis and Elisa Mandato PhD (Dana-Farber Cancer Institute) and Antonio Ferreira PhD (Brigham and Women’s Hospital) for discussions regarding site mutagenesis. The authors thank the Joslin Islet Isolation Core, Joslin Flow Cytometry Core, and Joslin Bioinformatics Core (P30 DK36836). This work is supported by NIH grants R01 DK67536 (R.N.K.), UC4 DK116278 (R.N.K. and C.H.), RM1 HG008935 (C.H.), and R01 DK122160 (W.J.Q. and R.N.K.). Portions of the mass spectrometry work were performed in the Environmental Molecular Sciences Laboratory, Pacific Northwest National Laboratory, a national scientific user facility sponsored by the Department of Energy under Contract DE-AC05-76RL0 1830. R.N.K. acknowledges support from the Margaret A. Congleton Endowed Chair and C.H. is a Howard Hughes Medical Institute Investigator. DFDJ acknowledges support by Mary K. Iacocca Junior Postdoctoral Fellowship and American Diabetes Association grant #7-21-PDF-140. The authors sincerely thank the families of the human islet donors.

## AUTHOR CONTRIBUTIONS

D.F.D.J. conceived the study, designed and performed experiments, analyzed the data, assembled figures, and wrote the manuscript. Z.Z. performed RNA-seq, m^6^A-seq, and m^6^A LC-MS/MS experiments. N.K.B. performed morphometric analyses of pancreases and assisted with animal experiments. X.L. and M.J.G. performed LC-MS/MS on S-nitrosylation. S.K. performed cell culture experiments. J.W. assistant with omics data handling. J.H. performed immunohistochemistry. G.B. Assisted with *in vivo* experiments. L.X. assisted with FACS experiments. T.M.R. provided reagents. C.M. provided microarray data on T1D islets. A.C.P. provided T1D islets. M.A.A. contributed to conceptual discussions and assisted with NPOD pancreatic sections. D.L.E. contributed to conceptual discussion and shared protocols. S.D.P. assisted with METTL3 structural modeling. A.P. assisted with human pseudoislet experiments. W.J.Q. performed LC-MS/MS data analysis and contributed to conceptual discussions. C.H. contributed to conceptual discussions, designed the experiments, and wrote the manuscript. R.N.K. conceived the study, designed the experiments, supervised the project, and wrote the manuscript. All the authors have reviewed, commented on, and edited the manuscript.

## DECLARATION OF INTERESTS

R.N.K is on the scientific advisory board of Novo Nordisk, Biomea, and Inversago. C.H. is a scientific founder and a member of the scientific advisory board of Accent Therapeutics. The remaining authors have no conflicts of interest.

## METHODS

### Human islet isolation and processing

Human islets were obtained from the Integrated Islet Distribution Program (IIDP), Prodo Laboratories, ADI isletcore, and provided by Alvin C Powers MD (Vanderbilt U). Freshly isolated islets were cultured overnight (16h) in Miami Media #1A (Cellgro) upon arrival. Islets were then handpicked, and seeded on ultra-low-attachment 6-well plates (Corning) (200 IEQ/well) for experiments. Snap-frozen control and T1D islets from ADI isletcore were immediately lysed in Trizol (ThermoFisher) upon arrival and stored at -80C for RNA isolation.

### EndoC-βH1 Cell Culture

The EndoC-βH1 cell line was cultured and passaged as previously described (Ravassard et al., 2011). Briefly, culture plates were coated with DMEM (glucose 4.5 g/L) containing PS (1%), fibronectin (2 µg/mL), and extracellular matrix (1% vol/vol) (MiliporeSigma) and incubated for at least 1 h in 5% CO2 at 37°C before the cells were seeded. EndoC-βH1 cells were grown on Matrigel/fibronectin-coated (MiliporeSigma) culture plates containing DMEM (glucose 1 g/L), BSA fraction V (2% wt/vol) (Roche Diagnostics), 2-mercaptoethanol (50 µM), nicotinamide (10 mM), transferrin (5.5 µg/mL), sodium selenite (6.7 ng/mL), and PS (1%) (Sigma-Aldrich) (Ravassard *et al.,* 2011).

### Human islets and EndoC-βH1 cell treatments

#### Cytokines treatments

Overnight cultured human islets or EndoC-βH1 cells were challenged with vehicle (PBS), IL-1 β (50 U/ml; R&D Systems, USA), IFN-α (2000 U/ml; PBL Assay Science), IFN-γ (1000 U/ml; Peprotech), or a combination of IL-1 β + IFN-α, or IL-1 β + IFN-γ in respective culture media. After treatments, islets were then handpicked, washed twice with ice-cold DPBS (GIBCO) by self-sedimentation, and immediately lysed in Trizol for RNA isolation, RIPA buffer (ThermoFisher), or fixed and embedded in agar for immunofluorescence staining as previously described (El Ouaamari et al., 2016).

#### Thapsigargin treatments

Overnight cultured human islets were treated with thapsigargin (1 µM; Selleckchem) or vehicle (DMSO) in Miami Media #1A for 16h and collected for protein isolation.

#### H_2_O_2_ and N-acetyl cysteine (NAC) treatments

Overnight cultured human islets were treated with 25 µM of H_2_O_2_ (MiliporeSigma), or H_2_O_2_ plus 1 mM of NAC (Cayman chemical), or vehicle (PBS) in Miami Media #1A for 24h and collected for protein isolation.

#### S-Nitroso-N-Acetyl-D,L-Penicillamine (SNAP) treatments

Overnight cultured human islets were treated with 10, 100, or 1000 nM of SNAP (Cayman chemical), or vehicle (PBS, pH 7.2) in Miami Media #1A for 24h and collected for protein isolation.

#### Actinomycin D treatments

EndoC-βH1 cells cultured as described above and at 48h post-seeding/knock-down or overexpression were challenged with IL-1 β + IFN-α as described above. At 72h post-seeding/knock-down or overexpression, cells were treated with 10µg/µL Actinomycin D (ThermoFisher) or DMSO for 0, 4, or 8h.

### Transfections

#### Knock-down experiments

Reverse transfections were performed as previously described (De Jesus *et al.,* 2019). Briefly, EndoC-βH1 cells or dispersed human islet cells were mixed with Lipofectamine RNAiMAX Reagent (Life Technologies) and small interfering RNA complexes (Dharmacon) at a final concentration of 15 nmol/L siRNA according to manufacturer instructions. EndoC-βH1 cells were seeded at a density of 6×10^4^ cells/cm^2^ in Matrigel/fibronectin-coated (MiliporeSigma) culture plates. Human dispersed islets were seeded at a density of 5×10^4^ cells/cm^2^ on ultra-low attachment plates (ThermoFIsher) and allowed to form spontaneous pseudoislets. EndoC-βH1 cells and human pseudoislets were collected 96h post-transfection. ON-TARGETplus Non-Targeting Control Pool D-001810-10-05, ON-TARGETplus METTL3 siRNA L-005170-02-0005, ON-TARGETplus Human YTHDF1 siRNA L-018095-02-0005, ON-TARGETplus Human YTHDF2 siRNA L-021009-02-0005, ON-TARGETplus Human YTHDF3 siRNA L-017080-01-0005 (Dharmacon, USA).

#### Overexpression experiments

EndoC-βH1 cells were seeded at a density of 6×10^4^ cells/cm^2^ in Matrigel/fibronectin-coated (MiliporeSigma) culture plates. After 24h media was changed and cells were forward-transfected with c-Flag pcDNA3 (addgene #20011) or pcDNA3/Flag-METTL3 (addgene #53739) using Lipofectamine 3000 (Invitrogen) and Opti-MEM (Invitrogen) according to manufacturer protocols. Media was exchanged after 8h of transfection, and at 48h cells were further used for experiments including cytokine treatments and Actinomycin D treatments.

### Mouse studies

Female NOD/shiLTJ (“NOD”; Jackson Laboratories #001976) and NOD.Cg-Prkdcscid Il2rgtm1Wjl/SzJ (“NOD NSG”; Jackson Laboratories #005557) mice were used. Mice were housed on a 12-h light/12-h dark cycle with water and food ad libitum. Female mice were used for all experiments throughout the study. Blood glucose was measured weekly for follow-up studies and mice were considered diabetic when two consecutive measurements of blood glucose exceed 250 mg/dL. Serum C-peptide levels were measured using ELISA kits (Crystal Chem) according to manufacturer guidelines All mice were kept in a specific pathogen-free facility in the Animal Facility at Joslin Diabetes Center, and animal protocols were approved by the Institutional Animal Care and Use Committee (IACUC). Sample sizes for animal experiments were chosen based on experience in previous in-house studies of metabolic phenotypes and to balance the ability to detect significant differences with minimization of the number of animals used following NIH guidelines.

#### Mouse Islet Isolations

Islets were isolated from female NSG mice as previously described (El Ouaamari et al., 2015). In brief, 3-month-old mice were anesthetized, and their pancreas was infused with liberase (Roche). Following incubation at 37°C for 17 min the digested pancreases were washed, filtered through a 400μm filter, and run on a Histopaque (Sigma, USA) gradient. The purified islets were handpicked, counted and cultured overnight in 7mM glucose RPMI media (Gibco, USA) containing 10% FBS and 1% PS) (Gibco, USA). Islets were handpicked, washed 2 times with ice-cold DPBS by self-sedimentation, and used for experiments.

#### β-cell sorting by FACS

Overnight cultured mouse islets were dispersed with a solution of 1 mg/ml trypsin and 30 µg/ml DNase followed by incubation for 15 min at 37°C. During the digestion, the islets were vortexed every 5 min for 10s. Cold media including serum was added to stop the digestion, and the cells were washed two times in DPBS containing 1% fatty-acid-free bovine serum albumin (BSA). Before sorting islet cells were filtered through a 35 µm filter and sorted using MoFlo Cytometer (Dako), where cells were gated according to forward scatter and then sorted based on endogenous fluorescence (Smelt *et al.,* 2008) and CD45 staining (Biolegend #QA17A26).

#### Co-culture of splenocytes and islets

Total splenocytes were purified as previously described (Dirice *et al.,* 2019). Briefly, freshly harvested spleens of 12-13 weeks-old female NOD mice with early diabetes were filtered through a nylon mesh by followed lysis of the red blood cells with ACK Lysing buffer (Lonza). After starvation, 100 size-matched islets from NOD NSG mice were co-cultured with NOD splenocytes in 5 mmol/L glucose RPMI at a ratio of 1:10 as previously described (Dirice et al., 2014). At 48h islets were hand-picked washed in ice-cold DPBS and lysed in trizol for RNA isolation.

#### *In vivo* Mettl3 overexpression

Adeno-associated virus serotype 8 (AAV8) overexpressing Mettl3 (NM_019721.2) or eGFP under the control of rat insulin II promoter (addgene # 15029) with a WPRE element were synthesized by VectorBuilder. Briefly, 4-week-old NOD female mice received an intraperitoneal injection of 200 µl PBS containing 1×10^11^ gene copies (gc)/mouse of AAV8 overexpressing eGFP or Mettl3 and were followed for 20 weeks.

### RNA isolation and RT-PCRs

Total RNA was isolated as previously described (De Jesus et al., 2020). In brief, high-quality total RNA (>200nt) was extracted using standard Trizol reagent (Invitrogen) according to manufacturer instructions and the resultant aqueous phase was mixed (1:1) with 70% RNA-free ethanol and added to Qiagen Rneasy mini kit columns (Qiagen) and the kit protocol was followed. RNA quality and quantity were analyzed using Nanodrop 1000 and used for reverse transcription using the high-capacity cDNA synthesis kit (Applied Biosciences). cDNA was analyzed using the ABI 7900HT system (Applied Biosciences) and gene expression was calculated using the “Ct method. Data were normalized to GADPH.

### Protein isolation and Western-blotting

Total proteins were harvested from tissue and cell lines lysates using M-PER protein extraction reagent (Thermo Fisher) respectively supplemented with proteinase and phosphatase inhibitors (Sigma) according to standard protocol. Protein concentrations were determined using the BCA standard protocol followed by the standard western immunoblotting protocol of proteins. The blots were developed using chemiluminescent substrate ECL (ThermoFisher) and quantified using Image studio Lite Ver. 5.2 software (LICOR).

### Pancreas immunostaining and analyses

Mouse pancreas was collected and fixed in 4% formaldehyde at 4°C overnight, followed by paraffin embedding. Five-micron-thick slides were cut and subjected to immunostaining. Slides were heated in 10mM sodium citrate, followed by blocking with donkey serum, and incubated with various primary antibodies: Insulin (Abcam #ab7842, Guinea pig polyclonal), Glucagon (Sigma #G2654, mouse monoclonal), Somatostatin (Abcam #ab64053, rabbit polyclonal). Specific signals were detected by using fluorescence-conjugated secondary antibodies (Jackson Immunoresearch, Alexa 488, Alexa 594, and AMCA). Images were captured using Zeiss Axio Imager A2 upright fluorescence microscope. Insulitis was evaluated as reported previously (Dirice *et al.,* 2019). Quantification of β-cell mass was performed as previously described (El Ouaamari *et al.,* 2016).

### Measuring total m^6^A levels

Total m^6^A levels were measured by employing LC-MS/MS or a quantitative colorimetric ELISA.

#### LC-MS/MS quantification of m^6^A

Total m^6^A levels among all adenosines were measured by triple-quad LC-MS/MS. We first purified mRNA from human islets total RNA by two rounds of polyA selection using polyA beads. 50 ng of purified mRNA were subject to digestion by 1 Unit of nuclease P1 (Sigma #N8630-1VL) in 25 µL of buffer containing 20 mM of NH4Ac at 42 degrees for 2 hours followed by phosphatase treatment using 1 µL of FastAP Thermosensitive Alkaline Phosphatase (ThermoFisher #EF0651) at 37 degrees for 1 hour. The digested nucleotides were filtered by a 0.22 µm syringe filter (Millipore) and then analyzed by a C18 reverse phase column on HPLC (Agilent) followed by triple quad MS/MS quantification (Sciex). The concentration of each type of nucleotide was calibrated by standard curves measured from pure nucleoside standards in each experiment. The m^6^A/A ratio was computed using the estimated m^6^A and A-concentrations.

#### Colorimetric quantification of m^6^A

EpiQuik m^6^A RNA Methylation Quantification Kit (EpigenTek) was used to measure the percentage of m^6^A methylation level in total RNA in EndoC-βH1 cells harboring WT or mutant METTL3 overexpression according to the protocols of the manufacturer using the kit provided negative control, positive control, and our samples consisting of 200µg of total RNA from EndoC-βH1 cells. The m^6^A percentage in total RNA was calculated using the following formula: m^6^A% = (Sample OD - NC OD) * S)/(PC OD - NC OD) * p) × 100%. NC: negative control; PC: positive control; S: the amount of input sample RNA; p: the amount of input positive control. Equal amounts of RNA samples were used.

### m^6^A immunoprecipitation and sequencing

For patient islets samples, polyA-selected mRNA was adjusted to 15 ng/µL in 100ul and fragmented using Bioruptor ultrasonicator (Diagenode) with 30s on/off for 30 cycles. m^6^A-immunoprecipitation (m^6^A-IP) were performed using the monoclonal m^6^A antibody from the EpiMark N6-Methyladenosine enrichment kit (NEB cat. E1610S). Input and eluted total RNA from m^6^A-IP were used to prepare libraries with Takara Pico-Input Strand-Specific Total RNA-seq for Illumina v2 (Takara). Sequencing was performed on Illumina Nova-seq according to the manufacturer’s instructions. Approximately 30 million paired-end 150-bp reads were generated for each sample.

### Differential methylation analysis for m^6^A sequencing

Human genome sequences and gene annotations were downloaded from the UCSC golden path, version hg38. We generated genome indexes using the genomeGenerate module of the STAR aligner (Dobin et al., 2013) with sjdbOverhang as 52 for 53-bp reads and 59 for 60-bp reads. The reads were trimmed for adapters and poly(A/T) tails, and then filtered by sequencing Phred quality (>= Q15) using fastp (Chen et al., 2018b). We aligned the adapter-trimmed reads to the genome using STAR with the two-pass option and indexed the BAM files with samtools (Li et al., 2009a). Using the R package RADAR (Zhang et al., 2019), we counted the mapped reads in 50-bp consecutive bins of each gene for each pair of input and m^6^A immunoprecipitation (IP) samples. Counts were normalized for library size and IP counts were adjusted for expression level by the gene-level read counts of input libraries. Bins with average IP-adjusted counts lower than 10 in both CTRL and CASE groups were removed. Then bins that were not enriched in IP were also filtered out. To construct PCA plots, we used the removeBatchEffect function in the limma package (Ritchie et al., 2015) to remove the subject effect for the human islets; the clone effect for the EndoC-βH1 cells; the batch, gender, age, and BMI effects for the T1D islet study. We perform differential methylation analysis of count data using the R package DESeq2 (Love et al., 2014). To consider the pairing of m^6^A IP and input, we use the normalized, expression-level-(i.e. input)-adjusted, and low-read-count-filtered IP counts. Using Wald test, we tested for significant effects of cytokine treatment or T1D on the m^6^A enrichment/depletion. To discover cytokine treatment effects in human islets or EndoC-βH1 cells, we performed paired tests so that each cytokine-treated sample was compared to its own paired baseline sample. To discover T1D effects in human islets using all controls, we adjusted for batch, gender, age, and BMI. We then merged the neighboring significant bins. P-values of these bins were combined by Fisher’s method (Fisher, 1992). We adjusted for multiple testing using the Benjamini-Hochberg false discovery rate (FDR) controlling procedure.

### Differential expression analysis of m^6^A-sequencing input samples

The input libraries of m^6^A sequencing are essentially mRNA sequencing libraries, so we performed gene-level differential expression analysis on them. After STAR alignment, alignments were assigned to genomic features (e.g. the exons for spliced RNAs) using featureCounts (Liao et al., 2014). Multi-mapping reads were counted as fractions. R package DESeq2 (Love *et al.,* 2014) was used to test for differential expression where sequencing batch, gender, and age were included as covariates. We adjusted for multiple testing using the Benjamini-Hochberg FDR procedure.

### RNA-seq analyses of sorted NOD mouse β-cells

Reversely stranded 100 bp single-end reads were trimmed for adapters and filtered by sequencing Phred quality (>= Q15) using fastp (Chen *et al.,* 2018b). Reads were aligned to the mouse transcriptome (Ensembl version 98) using kallisto (Bray et al., 2016) and transcript counts were converted to gene counts using tximport (Soneson et al., 2015). To filter out low-expressing genes, we only kept genes that had counts per million (CPM) more than 1 in at least 3 samples. We then normalized counts by weighted trimmed mean of M-values (TMM) (Robinson and Oshlack, 2010). To use linear models in the following analysis, we transformed counts into logCPM with Voom (Law et al., 2014). To discover the differential genes, we used the linear regression modeling R package limma (Ritchie *et al.,* 2015), which applied moderated t-tests to detect genes that are differentially expressed between groups. We adjusted for multiple testing using the Benjamini-Hochberg FDR procedure.

### Re-analyses of single-cell RNA-seq dataset

Data was obtained from GEO under the accession number GSE121863 (Russell *et al.,* 2019). First, we computed some quality control metrics with R package scater (McCarthy et al., 2017). We removed outlier cells whose library size and number of expressed genes were too low or whose proportion of counts assigned to mitochondrial genes was too high using thresholds of 2000, 1000, and 10%. We removed genes that have average counts of 0. We clustered similar cells together using a graph-based clustering algorithm and using genes that have average counts of more than 0.1. The algorithm normalizes the cells in each cluster using the deconvolution method (Lun et al., 2016). Finally, it performs scaling to ensure that size factors of cells in different clusters are comparable. Next, we estimated the technical noise by assuming the noise follows a Poisson distribution. We used the 1,000 genes that have the largest biological variations and performed Principal Component Analysis (PCA). We selected the first 100 PCs for the following analysis. We constructed tSNE plots from the PCs where each point represents a cell and was colored according to the variable diabetes. The insulin expression levels had 4 modes. We then used the Gaussian finite mixture model to identify cells in the 3rd and 4th mode, i.e. those with the highest insulin expression (Scrucca et al., 2016). To perform differential gene expression in β-cells, we selected β-cells and analyzed genes that are expressed in at least 10 cells using limma (Ritchie *et al.,* 2015), which applied moderated t-test to detect genes that are differentially expressed between the established T1D and Non-T1D. We adjusted for multiple testing using the Benjamini-Hochberg FDR procedure.

### Pathway enrichment analysis

Pathway enrichment analyses were performed using ConsensusPathDB using default settings (Herwig et al., 2016). GO terms tree were constructed using Cytoscape (Saito et al., 2012). Protein-protein functional networks were constructed using STRING using default settings (Snel et al., 2005).

### Biotin Switch assay

We used the biotin switch method that converts –SNO into biotinylated groups using a detection kit (Cayman Chemical), to detect S-nitrosylation of METTL3 in EndoC-βH1 cells treated with IFN-α and IL-1β according to the protocols of the manufacturer.

### LC-MS/MS analysis on METTL3 S-nitrosylation (SNO)

10 µg recombinant human METTL3 was incubated with 1 mM DTT for 30 min at room temperature (RT), and buffer was exchanged into 30 µL of 50 mM HEPES (pH 7.4) containing 1 mM EDTA, 0.1 mM neocuproine and 0.05% SDS using 30k spin columns. METTL3 was incubated with and without 200 µM SNAP in dark at RT for 1 hr. Excessive SNAP was removed by buffer exchange with 50 mM HEPES (pH 7.4). Free thiols in all samples were blocked by 20 mM NEM at RT for 30 min. NEM was removed by washing with 50 mM NH_4_HCO_3_(pH 8) containing 1 mM EDTA, 0.1 mM neocuproine, and 8M urea. SNO modifications in proteins were reduced with 20 mM sodium ascorbate and alkylated by 20 mM iodoacetamide at RT for 1 hr. Proteins were then digested by trypsin (enzyme to protein ratio = 1: 20) overnight. Peptides were eluted by centrifugation in 50 mM NH_4_HCO, and the concentration of each sample was adjusted to 0.05 µg/µL for LC-MS/MS. LC-MS/MS analysis was conducted using a nanoAcquity UPLC system (Waters) coupled to a Q-Exactive Mass Spectrometer as previously described (Duan et al., 2020). MS/MS raw data were searched against the Uniprot FASTA file of Homo Sapiens using MS-GF+ algorithm. Dynamic modifications included the oxidation of methionine (15.9949 Da), NEM on cysteine (125.0477 Da), and iodoacetamide on cysteine (57.0215 Da).

### Site-directed mutagenesis

Site-directed mutagenesis was performed by Genscript. Briefly, vector pcDNA3/Flag-METTL3 (addgene #53739) was transformed and the plasmid was extracted with Axygen kit (Corning). Next, sequence verification and enzyme digestion was performed for vector verification. Plasmid was linearized by digestion with HindIII and XboI to obtain a ∼5.2 kb vector backbone. Gene fragments were prepared by using two specific pairs of primers with overlap for each construct. Gene fragment amplification was performed by two rounds of PCR to generate the target gene fragment with the designed point mutation using the WT as a template. PCR product was purified and cloned, ligated with a linearized vector with T4 ligase. Gene products were transformed in competent cells and single colonies were picked for screening and sequence verification.

### METTL3 protein structure modeling

Structure figures of METTL3 were generated using Pymol (Delano) and coordinates from AlphaFold accession number AF-Q86U44-F1-model_v4 highlighting the zinc finger cluster and catalytic domain inter-domain interactions (also shown in the predicted aligned error (PAE) grid in Figure S5E) (Jumper et al., 2021). The zinc finger cluster is shown in light-orange and the catalytic SET domain in light-blue ribbon format and the SAM cofactor in stick format. The majority of the active site, including the cofactor and substrate binding pockets, is shown in dotted surface format. The solution structure of the zinc finger cluster from PDB 5YZ9 (Huang *et al.,* 2019) is also shown in greater detail in ribbon format including the zinc atoms and coordinating cysteine residues implicated in S-nitrosylation.

### Study Approval

All animal experiments were conducted following the Association for Assessment and Accreditation of Laboratory Animal Care. All protocols were approved by the Institutional Animal Care and Use Committee of the Joslin Diabetes Center following NIH guidelines. All human studies and protocols used were approved by the Joslin Diabetes Center’s Committee on Human Studies (CHS#5-05). Formal consent from human islet donors was not required because samples were discarded islets from de-identified humans.

## SUPPLEMENTARY FIGURES

**Supplementary Figure 1: Immune and antigen-presenting pathways are upregulated in Mettl14 deficient C57BL/6NJ mouse β-cells (related to Figure 1).**

**(A)** Venn diagram of differentially expressed genes in Mettl14 KO β-cells compared to controls. **(B)** Pathway enrichment analyses of genes upregulated in Mettl14 KO β-cells compared to controls. **(C)** Induced network analyses of significantly upregulated genes in Mettl14 KO β-cells and involved in the immune system and antigen presentation. *H2Aa, Cd44,* and *Cd74* are genes involved in antigen presentation and with the immunogenetics of Type 1 diabetes and are shaded in red. (Controls, n=4 pools, 2 animals/pool; M14KO, n=4 pools, 4 animals/pool). All samples in each panel are biologically independent. Statistical analyses were performed using the Benjamini-Hochberg procedure and genes were filtered for FDR<0.10. Data were downloaded and reanalyzed from dataset GSE132306 (De Jesus *et al.,* 2019).

**Supplementary Figure 2: Pre-diabetic NOD β-cells show enrichment in pathways associated with Type 1 diabetes, antigen presentation, and innate immunity (related to Figure 1).**

**(A)** Representative FACS sorting strategy to deplete CD45 positive cells, and obtain an enriched β-cell population based on size, granularity, and autofluorescence (FITC) from NOD female mice. **(B)** PCA plot of RNA-seq samples of 4- or 8-week-old FACS sorted β-cells or non-β-cells. **(C)** Heat-map representation of islet identity genes, showing enrichment for β-cell identity genes in the β-cell fraction compared to non-β-cells. **(D)** Heat-map of represented genes associated with Type 1 diabetes and upregulated in pre-diabetic 8-week-old β-cells compared to 4-week-old. **(E)** Heat-map of represented genes associated with antigen presentation and upregulated in pre-diabetic 8-week-old β-cells compared to 4-week-old. **(F)** Heat-map of represented genes associated involved in innate immunity and upregulated in pre-diabetic 8-week-old β-cells compared to 4-week-old. All samples in each panel are biologically independent. Data representing n=3 pools of 3 mice/group. Heat maps represent clipped Z-scored log CPM. Statistical analyses were performed using the Benjamini-Hochberg procedure and genes were filtered for FDR<0.05.

**Supplementary Figure 3: scRNA-seq re-analyses of established T1D islets, β-cell upregulation of METTL3 in human T1D onset, and Mettl3 downregulation in β-cells with T1D progression in female NOD mice (related to Figure 1).**

**(A-B)** t-SNE representation of β-cells (A) (high insulin expression) or α-cells (B) (high glucagon expression) in control and established Type 1 diabetes (T1D) (GSE121863) (Russell *et al.,* 2019). **(C)** Schematic representation of the Image J pipeline used to quantify the METTL3 nuclear intensity in proinsulin positive area in human pancreatic sections from the Network for Pancreatic Organ Donors with Diabetes (NPOD). **(D)** Representative low magnification pictures of immunofluorescence staining of METTL3 (green) and Proinsulin (red) in pancreatic sections from human control and T1D onset showing a robust islet enriched upregulation of METTL3 at T1D onset. **(E)** Representative pictures of immunofluorescence staining of Mett3 (green), Insulin (blue), and Cd3 (red) in pancreatic sections from NOD male mice. Males without any histological pancreatic immune cell infiltration patterns. Scale bar=100µM.

**Supplementary Figure 4: Human islets and EndoC-βH1 cells present an extensive overlap in the innate immune response to IL-1 β and IFN-α (related to** Figures 3 and 4). **(A)** Venn diagram representation of the upregulated (red), downregulated (blue), and unchanged genes (black) in EndoC-βH1 cells treated with IL-1 β and IFN-α compared to PBS. Statistical analyses were performed using the Benjamini-Hochberg procedure and genes were filtered for FDR<0.05. **(B)** Pathway enrichment analyses of upregulated and downregulated genes in EndoC-βH1 cells treated with IL-1 β and IFN-α compared to PBS. *P*-values were calculated according to the hypergeometric test based on the number of physical entities present in both the predefined set and user-specified list of physical entities. **(C)** Venn diagram representation of the commonly upregulated (red), downregulated (blue), and unchanged genes (black) of the intersected genes in EndoC-βH1 and human islets cells treated with IL-13 and IFN-α compared to PBS. Statistical analyses were performed using the Benjamini-Hochberg procedure and genes were filtered for FDR<0.05. **(D)** Pathway enrichment analyses of commonly upregulated and downregulated genes in human islets and EndoC-βH1 cells treated with IL-1β and IFN-α compared to PBS. *P*-values were calculated according to the hypergeometric test based on the number of physical entities present in both the predefined set and user-specified list of physical entities. **(E-F)** Volcano-plot representation of differentially expressed genes in human islets (E) and EndoC-βH1 cells (F) treated with IL-15 and IFN-α compared to PBS. Innate immune genes are depicted in red and show a near absolute overlap between human islets and EndoC-βH1 cells. **(G)** Metagene of m^6^A enriched peaks in PBS (blue) or IL-1β + IFN-α-treated (red) EndoC-βH1 cells. **(H)** Enrichment for known m^6^A consensus motif RRACH. **(I)** Histogram of the distribution of differential m^6^A loci log_2_ fold changes from IL-1β plus IFN-α-treated versus PBS in EndoC-βH1 cells. Human islets: n=15 biologically independent samples. EndoC-βH1 cells: n=6 biologically independent samples. Statistical analyses were performed using the Benjamini-Hochberg procedure and genes were filtered for FDR<0.05.

**Supplementary Figure 5: Summary schemes of experimental approaches and METTL3 protein structural modeling (related to Figure 7). (A)** Summary scheme related to Figure 7A of the experimental approach for the western-blot analyses of indicated proteins in human islets cells treated with H_2_O_2_ or H_2_O_2_ plus N-acetyl cysteine (NAC) for 24h (n=3/group). **(B)** Summary scheme related to Figure 7C of the experimental approach for western-blot analyses of indicated proteins in human islets treated with DMSO, IL1-β plus IFN-α, or pre-treated with thapsigargin plus IL1-β + IFN-α for a total of 24h (n=3/group). **(C)** Summary scheme related to Figure 7E of the experimental approach for western-blot analyses of indicated proteins in human islets treated with PBS or represented doses of S-nitroso-N-acetyl-DL-penicillamine (SNAP) for 24h (n=3/group). **(D)** Summary scheme related to Figure 7H of the experimental approach for S-nitrosylation site detection by LC-MS/MS. **(E)** Predicted aligned error related to Figures 7J and K. The color at position (x, y) indicates AlphaFold’s expected position error at residue x, when the predicted and true structures are aligned on residue y. This is useful for assessing inter-domain accuracy. **(F)** Conceptual model on the role of METTL3 in controlling the antiviral innate immune response by regulating the formation of deleterious dsRNAs and/or controlling viral response.

