## Supplementary Figures for "Redox Regulation of m^6^A Methyltransferase METTL3 in Human β-cells Controls the Innate Immune Response in Type 1 Diabetes"

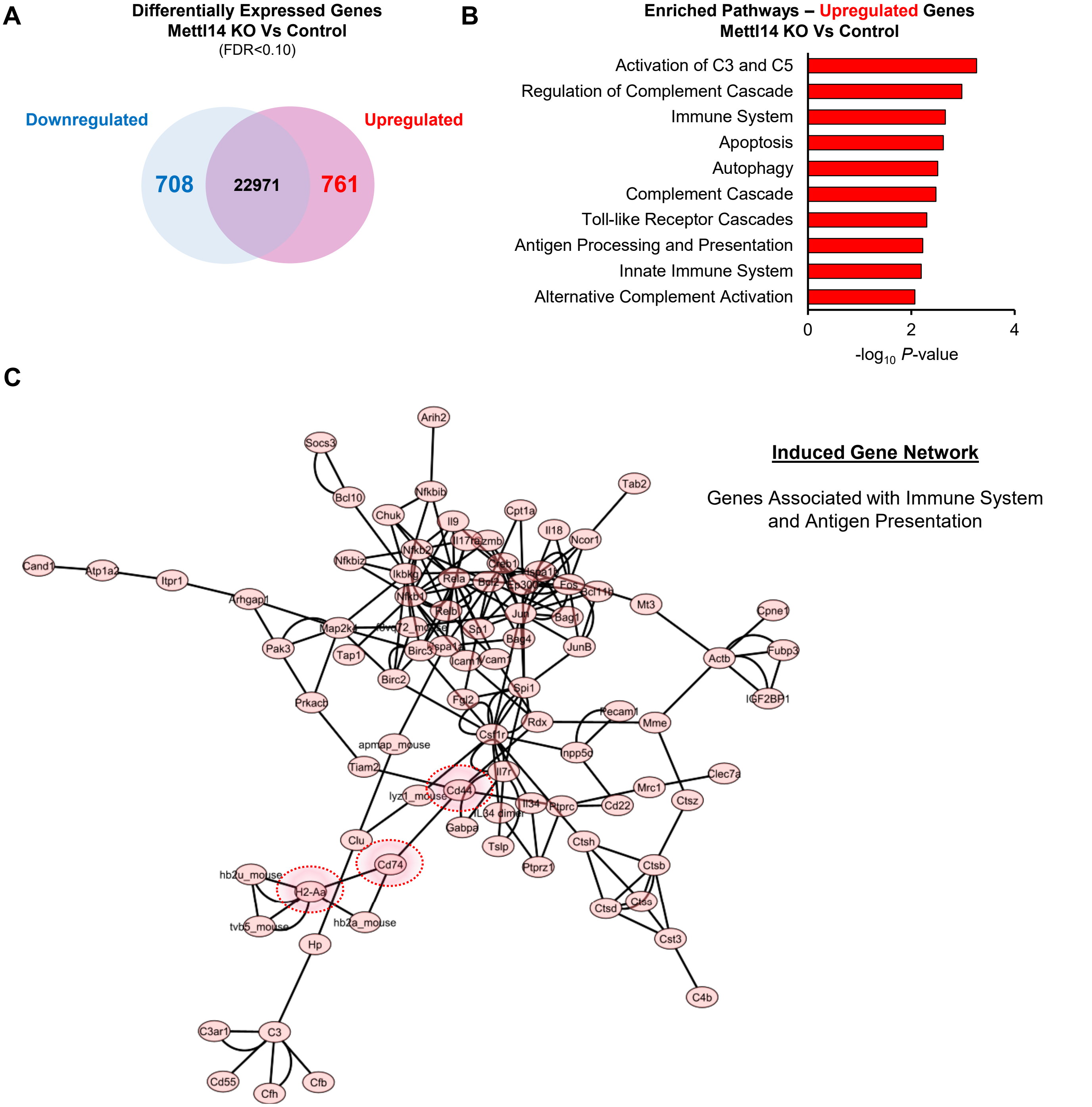


**SUPPLEMENTARY FIGURE 1**
**(Related to Figure 1)**

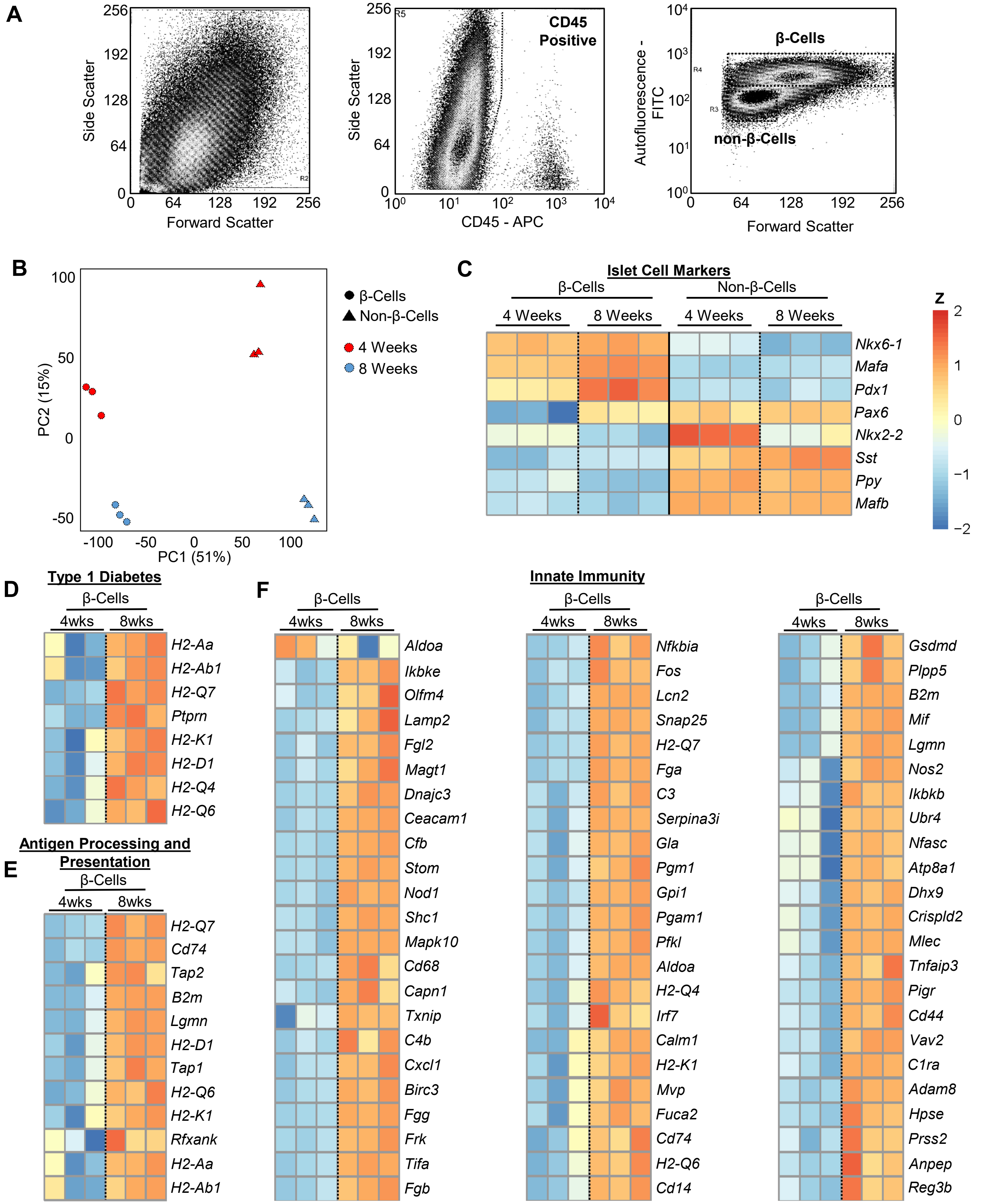


**SUPPLEMENTARY FIGURE 2**
**(Relative to Figure 1)**

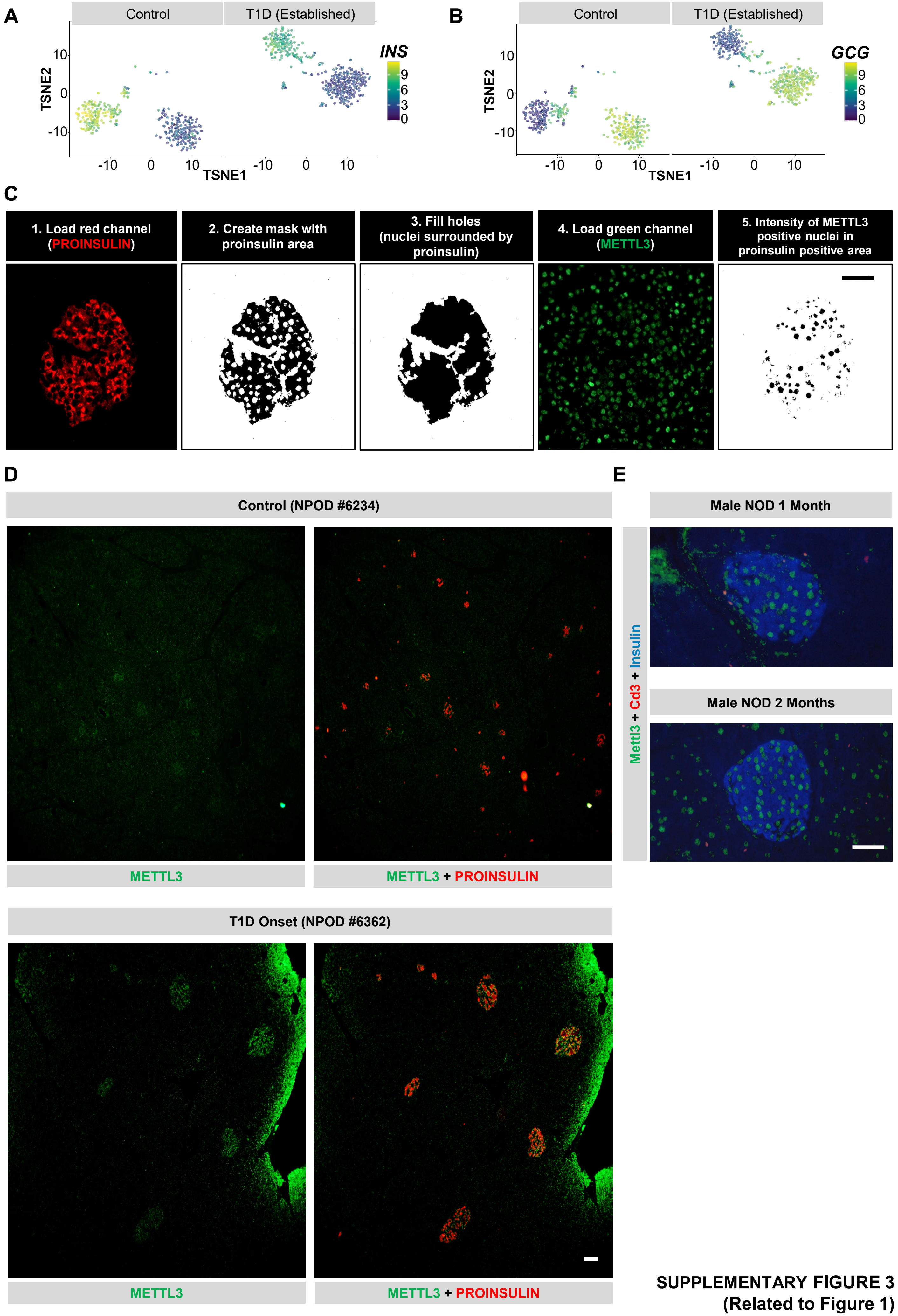


**SUPPLEMENTARY FIGURE 3**
**(Related to Figure 1)**

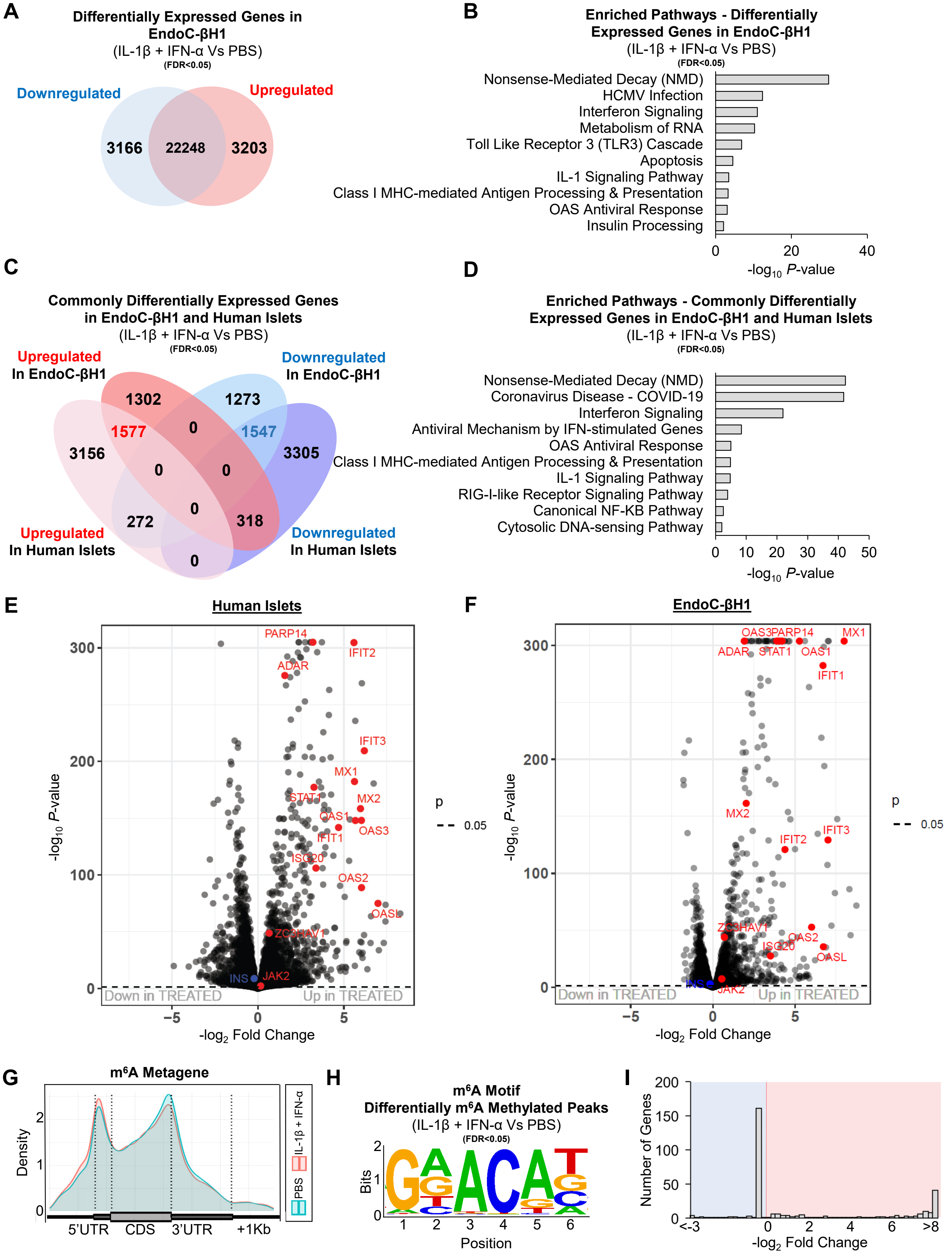


SUPPLEMENTARY FIGURE 4 (Related to Figure 3 and 4)

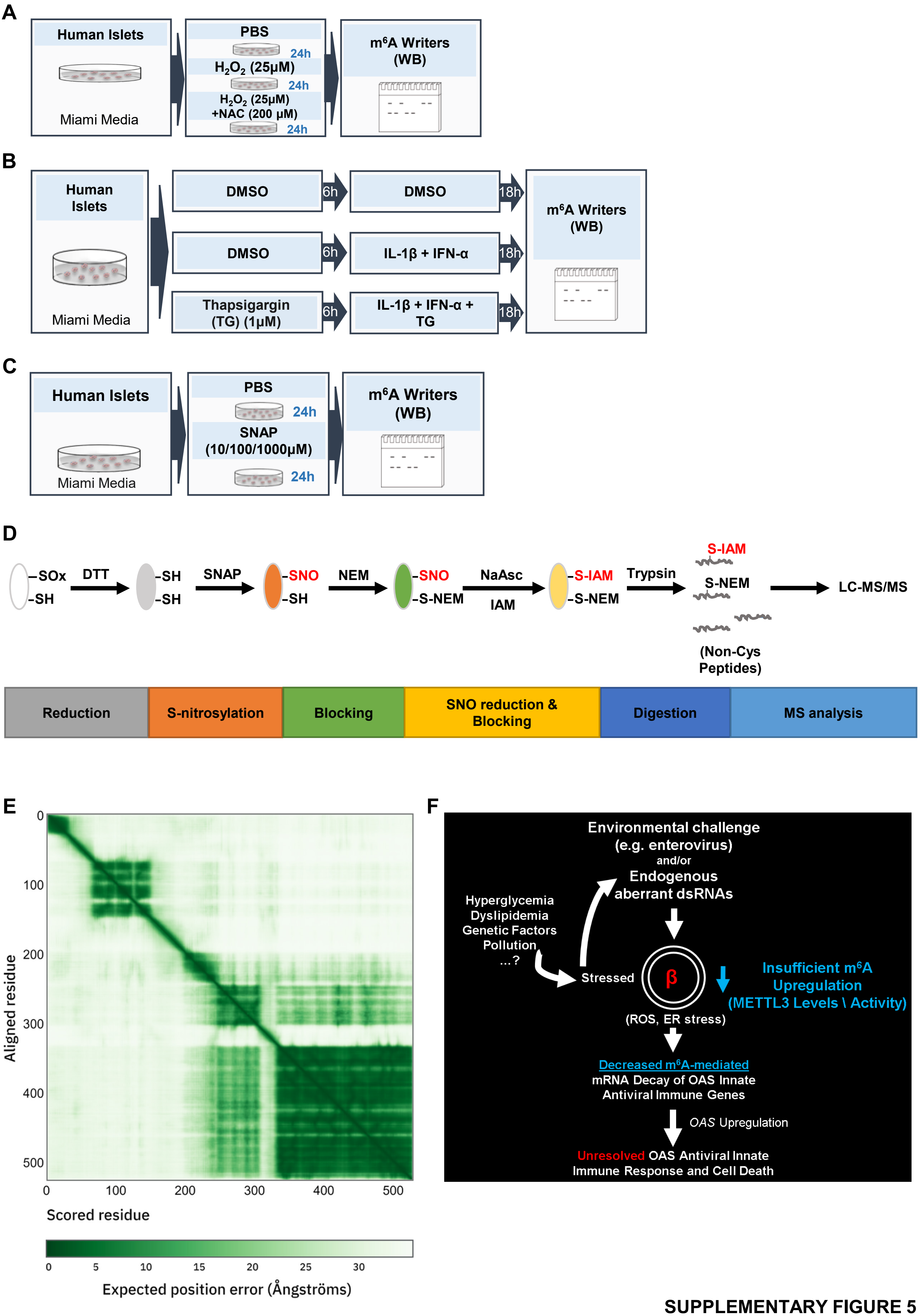


**SUPPLEMENTARY FIGURE 5**
**(Related to Figure 7)**
